# Pervasive polygenicity of complex traits inflates false positive rates in transcriptome-wide association studies

**DOI:** 10.1101/2023.10.17.562831

**Authors:** Yanyu Liang, Festus Nyasimi, Hae Kyung Im

## Abstract

Transcription-wide association studies (TWAS) and related methods (xWAS) have been widely adopted in genetic studies to understand molecular traits as mediators between genetic variation and disease. However, the effect of polygenicity on the validity of these mediator–trait association tests has largely been overlooked. Given the widespread polygenicity of complex traits, it is necessary to assess the validity and accuracy of these mediator–trait association tests. We found that for highly polygenic target traits, the standard test based on linear regression is inflated, leading to greatly increased false positives rates, especially in large sample sizes. Here, we show the extent of the inflation as a function of the underlying GWAS sample size and polygenic heritability of the target trait. To address this inflation, we propose an effective variance control method, similar to genomic control, but which allows for a different correction factor for each gene. Using simulated and real data, as well as theoretical derivations, we show that our method yields calibrated false positive rates, outperforming existing approaches. We further demonstrate that methods analogous to TWAS that associate genetic predictors of mediating traits with target traits suffer from similar inflation issues. We advise developers of genetic predictors for molecular traits (including polygenic risk scores, PRS) to compute and provide the necessary inflation parameters to ensure proper false positive control. Finally, we have updated our PrediXcan software package and resources to facilitate this correction for end users.

## Introduction

To explain the mechanisms behind the hundreds of thousands of loci discovered via genome-wide association studies (GWAS), researchers have studied the role of molecular traits as mediators. This has led to the development of transcriptome-wide association studies (TWAS) and related methods for other molecular traits (Gamazon et al., 2015; Gusev et al., 2016; Zhang et al., 2022). These mediator–trait association studies have been increasingly important in genetic studies.

TWAS and related methods, termed here xWAS, perform association tests between genetic components of molecular traits with the target trait, one feature at a time. There are known limitations of this approach that increase false positive rate because of molecular pleiotropy (i.e., when the same variant affects multiple genes but only one alters the target trait) and linkage disequilibrium (LD) contamination (i.e., when the variant altering the molecular trait has no effect on the trait but is in LD with a trait altering variant) (Wainberg et al., 2019; Zhu et al., 2016; Yuan et al., 2020; Zhao et al., 2024; Mancuso et al., 2019). Despite some of these limitations xWAS methods, including CWAS, RWAS, PWAS, and isoTWAS, are widely acknowledged as useful for nominating molecular traits driving the etiology of complex traits (Baca et al., 2022; Grishin and Gusev, 2022b; Zhang et al., 2022; Bhattacharya et al., 2023).

Some prior studies have reported potential inflation of false positives in TWAS. For example, van Iterson et al. (2017) argued that TWAS results tend to be biased and inflated, as indicated by deviations from the expected null distribution. The assumption that most features should not be associated with the target trait may be invalid due to the broad polygenicity of complex traits (Boyle et al., 2017). The authors concluded that the standard genomic control method overcorrected for this observed inflation and proposed a Bayesian approach to estimate the empirical null distribution as a solution. However, we demonstrate that this approach does not fully resolve the inflation caused by the target trait’s polygenicity.

de Leeuw et al. also suggested that TWAS may produce inflated type I error, attributing this inflation to inaccuracies in predicting gene expression traits (de Leeuw et al., 2023). However, error-in-variables theory (Fuller, 1987) assures us that noisy predictors reduce the power of the association, but does not cause inflation of type I error as long as the prediction error is not itself associated with the outcome. In line with this, our findings indicate that this inflation is better explained by the polygenicity of the target trait, rather than by prediction error itself.

It is increasingly accepted that there is widespread polygenicity of most complex traits (Yang et al., 2010; O’Connor et al., 2019; Boyle et al., 2017). The effect of polygenicity has been explored and leveraged in the context of GWAS with methods such as LD score regression and related approaches (Bulik-Sullivan et al., 2015). However, the effect of polygenicity on xWAS has not been explored rigorously.

In this study, we show that even in the absence of known false positives caused by molecular pleiotropy and LD contamination, the widespread polygenicity of the target trait—where most variants have a modest effect—leads to inflation of the association statistic. In other words, the false positive rate is higher than estimated by standard methods. To maintain the utility of xWAS methods, it is essential to ensure that false positive rates are well-calibrated.

We begin by demonstrating that polygenicity induces inflated type I error, even in a simple setup with independent SNPs and error terms. Next, we show this inflation in real data by assessing the association between genetically predicted expression in the UK Biobank and null polygenic traits. We further show that this inflation is not limited to a specific software but is a broader property of the TWAS/xWAS approach. Our analysis reveals a linear relationship between inflation, GWAS sample size, and the heritability of the target phenotype, which aligns with our theoretical derivations based on typical assumptions in statistical genetics. Finally, we propose a correction strategy, variance control, and demonstrate its effectiveness using both null and actual GWAS traits.

Finally, we modify the PrediXcan software to account for inflation and provide the necessary resources to implement this correction for expression predictors across 49 GTEx tissues (Barbeira et al., 2021), 1,156 metabolite predictors from the METSIM study (Yin et al., 2022), and 471 brain features used in the BrainXcan software (Liang et al., 2022).

## Results

TWAS and related methods nominate potential causal mediators (gene expression, protein levels, etc.) by testing the effect of the mediating trait ***T*** on a target trait ***Y***. We describe the model and usual assumptions here.

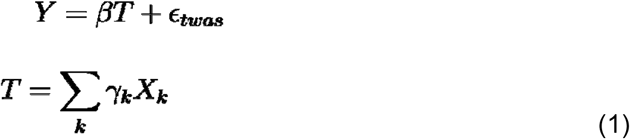

where *β* is the (fixed) effect of the mediating trait on the target trait, and ***ϵ***_***twas***_ is the error term independent of the mediator. For each genetic variant ***k***, the ***γ***_***k***_ is the genetic effect on the mediator, and the ***X***_***k***_ is the genotype dosage. This model accommodates both sparse (where most ***γ***_***k***_ = 0) and polygenic architecture (where most ***γ***_***k***_ **≠** 0) for the mediating trait. ***T*** here is the genetic component of the mediator.

### No inflation under ideal TWAS assumptions

To build intuition, we simulated target traits (***Y***) and unrelated mediating traits (***T***) under a simplified scenario. We refer to any target trait ***Y*** that is unrelated to the mediator ***T*** as a “null target trait.” Null target traits were simulated from a normal distribution for a sample of 1,000 individuals. For the mediating trait ***T***, we simulated 999 independent SNPs for the same 1,000 individuals and computed ***T*** as a weighted average of SNP dosages, with simulated genetic effects ***γ***_***k***_ drawn from a normal distribution. To ensure robustness to deviations from normality, we also repeated the simulation using t-distributed null traits and genetic effects. We performed this simulation 1000 times. For each simulation, we regressed ***Y*** on ***T*** and computed the Z-score, i.e., the ratio of the estimated effect of ***T*** on ***Y*** to its standard error.

A well-calibrated test should produce a Z-score that, under the null, approximately follows a standard normal distribution due to the central limit theorem of regression coefficients. Specifically, the sample variance of Z-scores obtained by regressing many null target traits (***Y***) on unrelated mediators (***T***) should approach 1 as the sample size increases. If the sample variance significantly exceeds 1, it indicates that the test is inflated.

As expected, we found that the association statistic is well-calibrated (Figure 1a-c), with p-values, Z-scores, and the sample variance of Z-scores following their expected distributions. Thus, when the standard TWAS assumptions hold, no inflation is observed.

**Figure 1:**
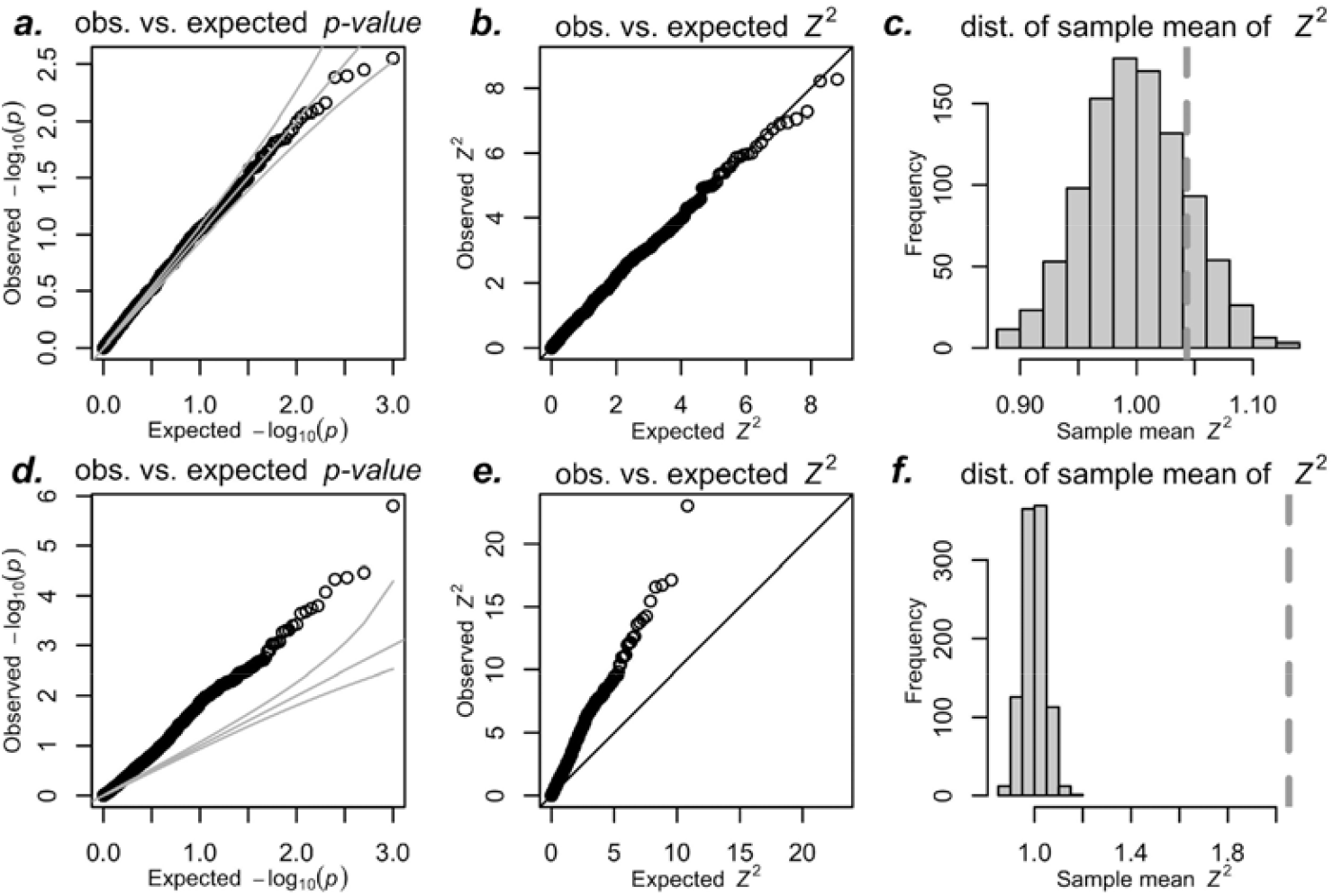
Inflation of simulated TWAS - simplified setting. This figure illustrates simulated TWAS under the null hypothesis using standard assumptions of independent error and mediator, following equation (1). We performed 1000 simulated associations with predicted expression using 1000 SNPs, with effect sizes and errors sampled from normal distributions. See Supplementary Figure S1 for simulations using Student’s *t*-distributed errors. The top row shows non-polygenic null trait simulations: (a) QQ-plot of observed p-values, (b) QQ-plot of observed *Z*^2^, and (c) average *Z*^2^ over 1000 simulations a dotted vertical line. These follow expected distributions under the null (uniform for p-values and standard χ with 1 degree of freedom). The average *Z* over 1000 simulations falls within the expected distribution (gray histogram in c) of sample means of standard χ^2^ with 1 degree of freedom. The bottom row present polygenic null trait simulations: (d) QQ-plot of observed p-values vs. expected, (e) QQ-plot of observed *Z*^2^ vs expected, and (f) average *Z*^2^ as dotted vertical line with a histogram of expected sample averages of squared standard normal random variables. All three panels d-f show departure from expected distributions under the null.

### No inflation with error in prediction independent of trait

In practice, we do not know the “true” weights for the genetic component of the mediator ***T***, meaning we use a set of weights 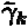 that differ from the true values ***γ***_***k***_. This includes cases where ***γ***_***k***_ is zero but 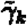 is not, and vice versa. Since prediction weights are typically trained in studies independent of the GWAS, it is reasonable to assume that the prediction error will be independent of the error term in the target trait ***Y*** (i.e., ***ϵ***_***twas***_ under the null). Under this assumption, the error-in-variable literature states that the association test using a nois explanatory variable remains valid, meaning there is no inflation of type I error (Fuller, 1987).

Intuitively, this result makes sense: under the null, when the mediator ***T*** is unrelated to the target trait ***Y***, adding error to ***T*** is unlikely to strengthen the association, provided the prediction error is independent of the target trait. If, however, prediction errors were systematically associated with target traits, TWAS results would become invalid, and the field would need to halt its use until a solution is developed. That said, most researchers would likely view this as an extreme measure and agree that, under the null, assuming independence between prediction error and the target trait is reasonable.

### Polygenic background of target trait causes inflated type I error

Next, we investigated whether a polygenic background in *Y* independent of the mediating trait inflates false positive rate (type I error) using the same simulation setting. We add a polygenic component to target trait by modeling the error term ***ϵ***_***twas***_ in equation (1) as follows:

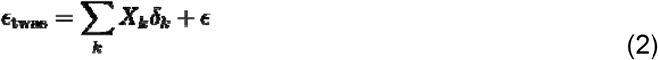

where ***δ***_***k***_ is sampled from a normal distribution with mean 0 and variance 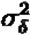. We also performed the simulation using t-distributions to check robustness to deviations from normality.

If this new component is independent of the mediator ***T***, the usual regression assumption holds, and hence, we did not expect inflated type I error, i.e., we expected that the variance of the Z-score statistic would have variance 1. However, contrary to our intuition, the sample variance of the Z-score statistic was much larger than its expected distribution as shown in Figure 1d-f.

### TWAS with real data also yields inflation when the target trait is polygenic

To test whether inflation occurs in TWAS using real data, we predicted the expression of *AMT*, chosen as a representative example, using genotype data from the UK Biobank along with prediction weights from Predictdb.org and the Fusion website (Gusev et al., 2016).

Calculating type I error rates requires operating under the null hypothesis, meaning we need traits that are unrelated to any genes. However, since all UK Biobank traits show significant GWAS loci, real traits could not serve this purpose. Instead, we generated simulated null traits. For non-polygenic null traits, we sampled values from a normal distribution. For polygenic null traits, we used UK Biobank genotype data, constructing each trait as a linear combination of genotype dosages with randomly generated weights from a normal distribution. To confirm robustness, we repeated the analysis using Student’s t-distributed weights, finding no significant change in results and thus demonstrating the robustness of our approach to deviations from normality.

For non-polygenic null traits, TWAS p-values, squared Z-scores, and sample variance of the Z-scores all follow expected distributions, showing no type I error inflation (Figure 2a-c). This outcome aligns with error-in-variable theory, which predicts that noisy predictors do not inflate type I errors when the noise is uncorrelated with the outcome (Fuller, 1987). These results confirm that noisy gene expression predictors do not cause inflation for non-polygenic traits.

**Figure 2:**
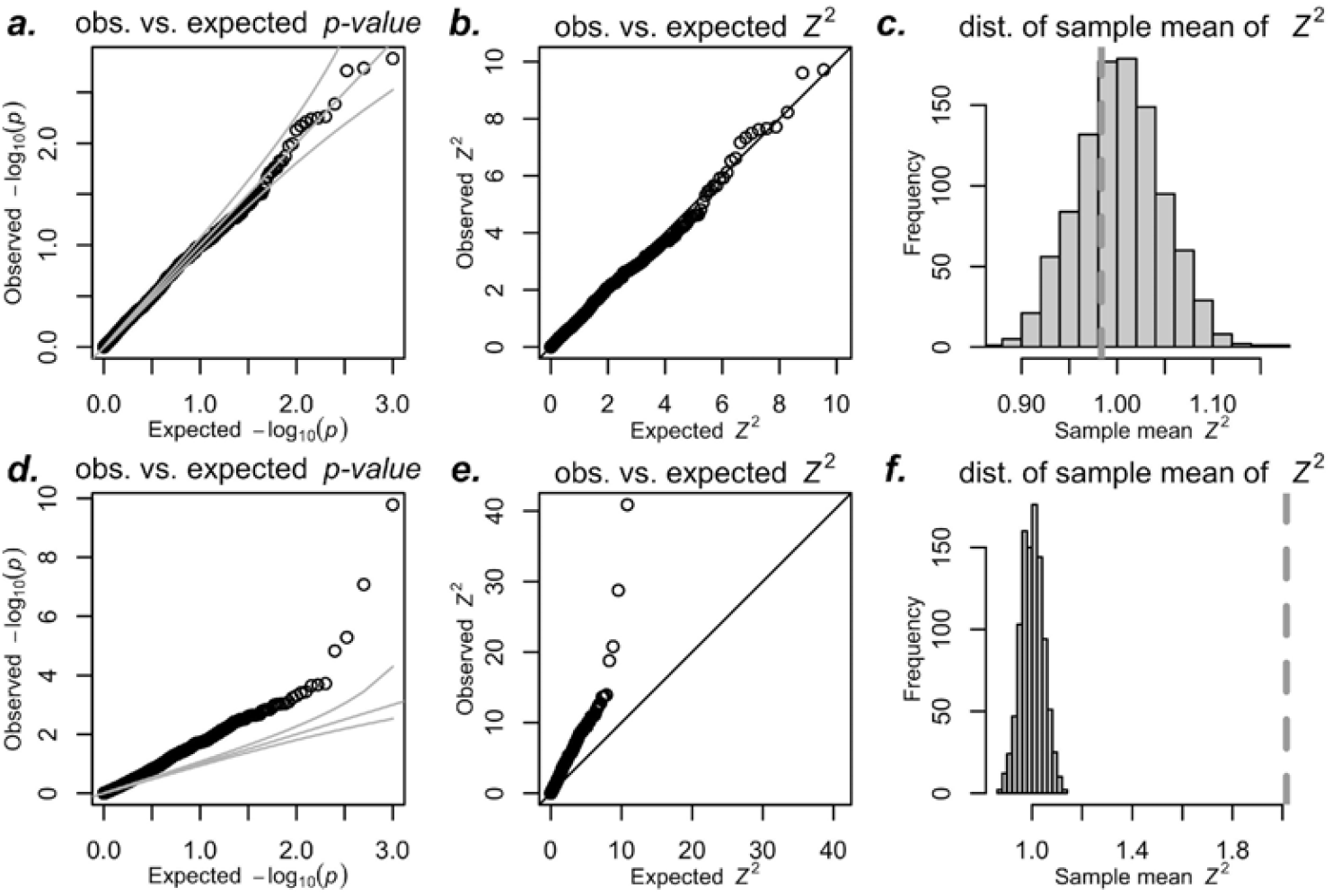
Inflation in real TWAS in the UK Biobank. We predicted expression of the gene *AMT* in whole blood for 10K randomly sampled white British unrelated individuals from the UK Biobank. For panels a-c, we simulated non-polygenic target traits from a normal distribution. For panels d-f, we simulated polygenic target traits as the sum of a polygenic component and independent normally distributed noise. We regressed the target trait on the predicted expression and calculated Z-scores, repeating this 1000 times for both trait types. Panels a and d show p-values, while b and e show squared Z-scores (*Z*^2^) for non-polygenic and polygenic traits, respectively. Panels c and f display the sample mean of *Z*^2^ as a vertical dotted line, with histograms showing sample means of squared standard normal variables to illustrate expected means under the null. Prediction weights for *AMT* were downloaded from PredictDB.org. Analogous figures using Fusion weights are shown in Supplementary Figure S3. Supplementary Figure S2 demonstrates the simulations with Student’s t distribution to show robustness to deviation from normality.

For polygenic null traits, however, we observed significant deviations from expected distributions in p-values, Z-scores, and sample variance of the Z-scores (Figure 2d-f). Moreover, similar inflation was observed using the Fusion software and prediction weights from the Fusion/TWAS website (Supplementary Figure S3), indicating that this issue is intrinsic to the TWAS/xWAS approach rather than specific to any particular implementation.

To further understand this inflation, we investigated how it varies with target trait heritability and GWAS sample size.

### Inflation grows linearly with the trait heritability and the GWAS sample size

We examined the relationship between the sample variance of the Z-score and both sample size and target trait heritability using 1000 simulated polygenic null traits in the UK Biobank. Figure 3 shows that the sample variance of the Z-score increases linearly with both factors. When the trait had no polygenic component (*h*^2^ = 0), the sample variance is approximately 1, as expected. However, the rate of inflation varies among genes, suggesting a straightforward formula for predicting the variance of the Z-score:

**Figure 3:**
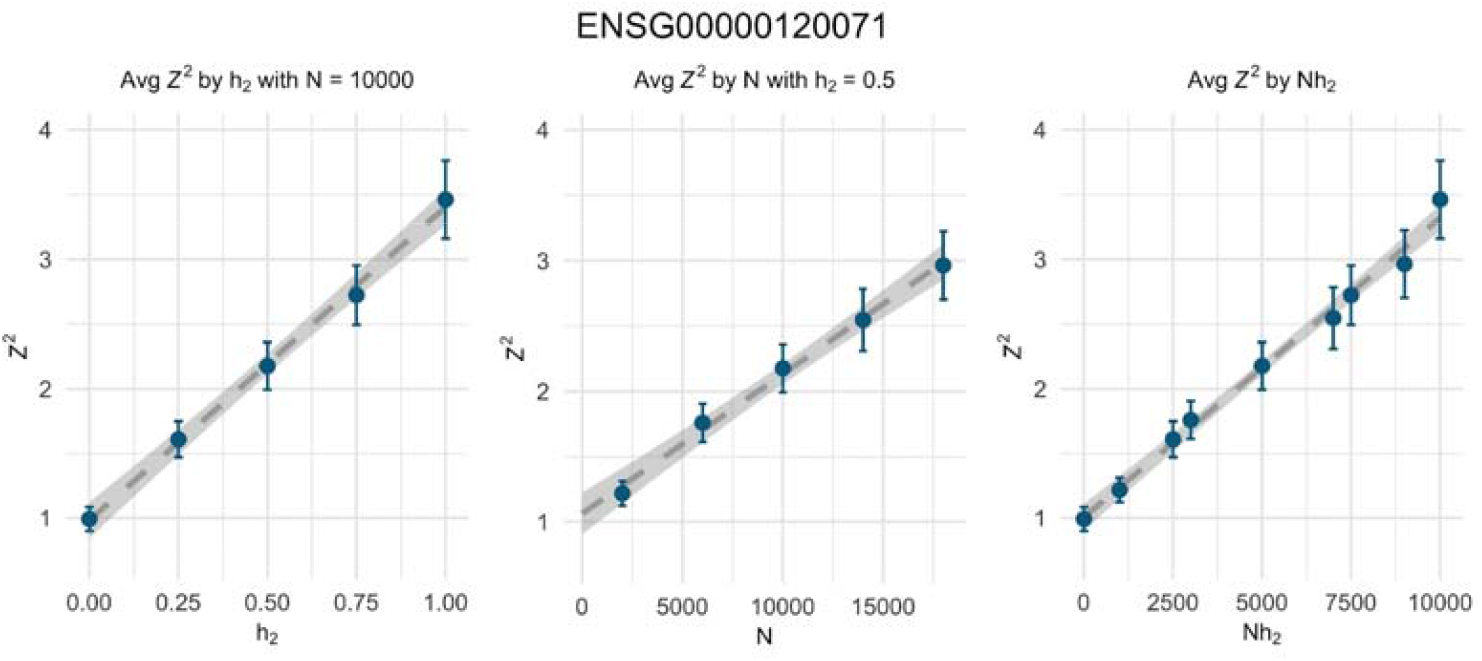
Linear dependence of inflation on the GWAS sample size and heritability of the target trait. The average *Z*^2^ is plotted against heritability 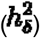, sample size (*N*), and the product of heritability and sample size 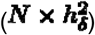. Each dot represents the average association of predicted expression of *KANSL1* with UK Biobank genotype data and 1000 null polygenic traits in the same individuals, at the specified sample sizes and heritability values. The error bars show 1.96x the standard errors of the sample averages. Dashed lines correspond to estimated linear regression lines based on the *Z*’s and the gray band represents the confidence interval of the regression lines. Supplementary Figures S4 and S5 illustrate similar trends using metabolite and brain image-derived features as mediators.

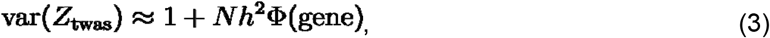

where Φ is the slope of inflation as a function of 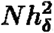.

In fact, our theoretical derivation, based on reasonable assumptions commonly made in statistical genetics (see Supplementary Note), demonstrates that var(*Z*_twas_) aligns with the formula in Eq.(3) with an “inflation slope parameter”

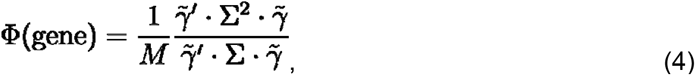

where Σ is the genome-wide LD matrix, *M* is the effective number of causal SNPs for the targ trait, and 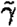 is the vector of prediction weights downloadable from various publicly available databases (e.g., Fusion, predictdb.org, omicspred.org).

Due to the challenges in accurately estimating the effective number of causal variants (*M*), the true LD matrix, and in validating key assumptions, we opted for an empirical approach. To estimate the inflation slope Φ, we conducted TWAS using multiple simulated null target traits within a reference panel, as described in the following section.

### Other xWAS have the same inflation issue

We tested this inflation using mediators with different genetic architectures, such as metabolite levels and MRI-derived brain features, and found that they exhibit similar inflation patterns, as suggested by our theoretical derivations. See Supplementary Figures S4 and S5.

### Theoretical derivation including non-zero mediating effect

We extended the theoretical formula for the variance of the Z-score statistic to account for cases where the mediator effect (***β***) is non-zero. Additionally, we modeled the prediction weights 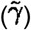 as the sum of the true weight (***γ***) and an independent error term, uncorrelated with both the true expression and the target trait. These assumptions are necessary to derive the equations under the alternative, but not under the null.

We demonstrated that the expected values of 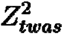 is given by

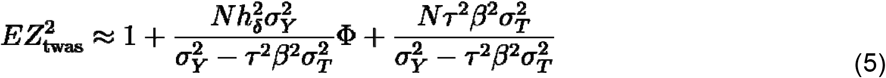

where *N* is the sample size, is the true effect of the mediator *T* on the target trait *Y*, and 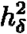 is the polygenic portion of *Y*, **Φ** is the inflation slope defined above in equation (4), and 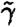 is the M-dimensional vector of noisy prediction weights, ***τ***^2^ is the precision of the prediction of the mediator. 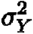 and 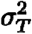 are the variances of the target (*Y*) and mediator (*T*) traits, respectively. See details in the Supplementary Note.

Precision of the prediction 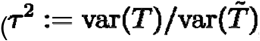, where 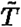 is the noisy version of the mediator) only appears in the formula alongside the mediator effect, specifically as ***βτ***^2^. (Note that the inflation slope parameter Φ does not depend on the precision, only on 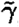, the LD matrix, and *M*.) This indicates that under the null hypothesis, where β = 0, prediction precision has no effect on type I error. This formula further emphasizes the fact that error-in-prediction does not cause inflation. Supplementary Figure S8a illustrates this independence of type I error from prediction precision. Panel b of the same figure shows how reduced prediction precision diminishes power; as lower Z-score variance decreases, the likelihood of achieving significance under the alternative hypothesis (when β ≠ 0).

### Variance control strategy to correct for the inflation

The p-values of TWAS associations are typically calculated under the assumption that under the null hypothesis the Z-scores follow a standard normal distribution, *N*(0,1). However, as demonstrated above, when target traits are fully polygenic, the variance of the Z-scores exceeds 1. This results in more extreme Z-score values and, consequently, more significant p-values than expected under the null hypothesis. To ensure reliable results, this inflation must be corrected.

Our approach is to adjust this inflation by dividing the Z-score by 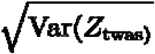, the square root of the Z-score’s variance. This variance increases linearly with both the GWAS sample size and the heritability of the target trait. Therefore, we propose estimating the slope of the inflation factor and using it to calculate a correction factor applicable to any sample size and heritability.

Once we determine the slope parameter Φ, it can be applied to GWAS of any size and target traits with polygenic architecture (Eqs. 1 and 2).

### Estimating the inflation factor

To estimate the slope parameter for each gene or mediator, we employed an empirical approach. For each predicted gene or mediating trait, we: (1) simulated a large number of null phenotypes *Y* across various sample sizes and heritability values; (2) performed association tests and calculated the *Z*^2^ values; and (3) averaged the *Z*^2^ values from each simulation. We then estimated the inflation slope **Φ** as the regression coefficient of average *Z*^2^ on the product of sample size and heritability.

We applied this estimation method to protein coding gene expression predictors in 49 tissues from GTEx (Barbeira et al., 2021), 1192 metabolite predictors trained with METSIM data (Yin et al., 2022), and 471 MRI-derived brain features in the UK Biobank (Liang et al., 2022).

The estimates of **Φ** varied across genes, metabolites, and brain features. Any negative estimates of **Φ** (genes n=34, metabolites n=34, brain features n=0) were set to zero as the lowest possible value. As shown in Figure 4, gene expression had the widest range of estimated **Φ** values (0 to 2 × 10^−4^, median 2 × 10^−5^), followed by metabolites (0 to 1.5 × 10^−4^, median 3.3 × 10^−5^), and brain features (3.9 × 10^−5^ to 6.5 × 10^−5^, median 5.3 × 10^−5^). Metabolites and brain features, which are more polygenic than gene expression, showed more consistent slopes. Most mediating traits (i.e., 78% of genes, 94% of metabolites, 100% of brain features) had **Φ** values in the 10^−5^ range.

**Figure 4:**
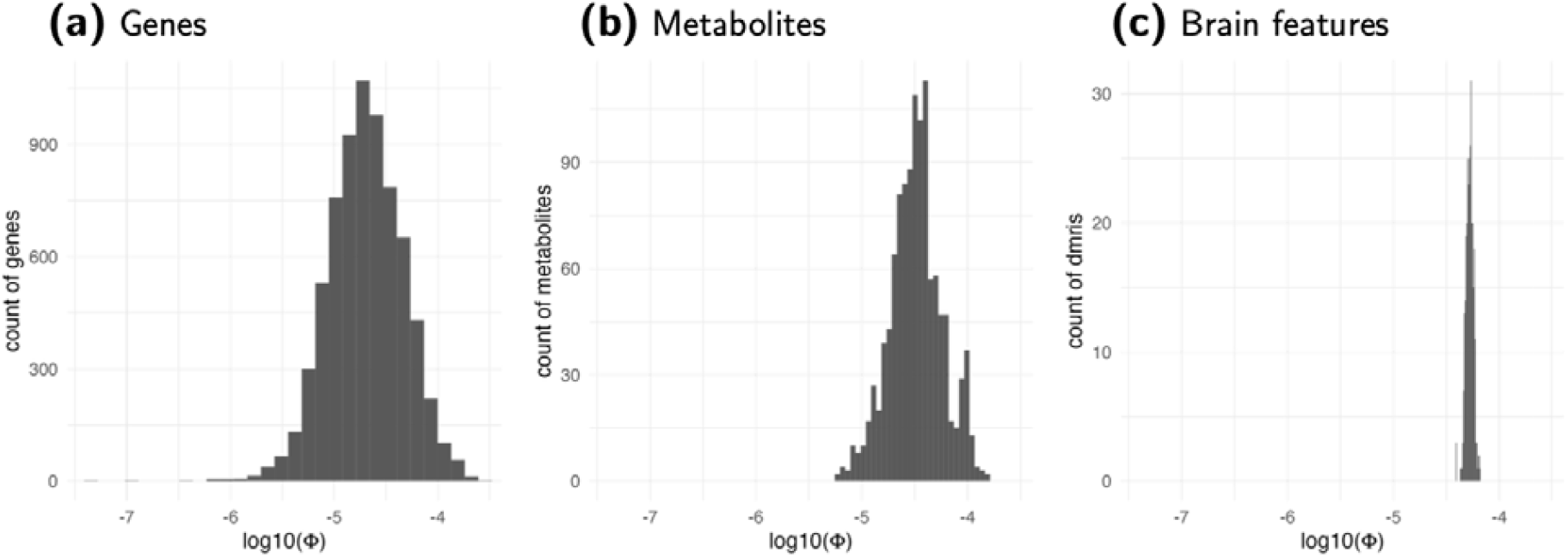
Distribution of Estimated Φ. Inflation factors for gene expression, metabolites, and brain features (diffusion MRI) are shown in the log10 scale. The factor for each mediator is estimated using the average Z^2^ statistics of the association between the genetically predicted mediator and 1,000 simulated target traits for each combination of heritability of target trait 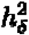 and sample size ***N***. The slope of the regression of 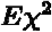 on 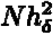 is used to estimate **Φ**. Most **Φ** values (78% of genes, 94% of metabolites, 100% of brain features) are in the range of 10^−5^.

### TWAS of null phenotype yields calibrated p-values after variance control correction

To demonstrate that our correction strategy provides calibrated false positive rates, we conducted TWAS on a null target trait in UK Biobank individuals using predicted whole blood gene expression. Rather than focusing on a single gene, we included all protein-coding genes with a single null target trait *Y* to recreate a TWAS in practice. Each gene was adjusted with its unique correction factor, based on the estimated **Φ**. As shown in Figure 5, our variance control method yields calibrated p-values (green crosses) that closely follow the expected line. By contrast, BACON, an alternative correction method (van Iterson et al., 2017), undercorrects inflation except in brain features, as indicated by the deviation of blue circles from the expected line.

**Figure 5:**
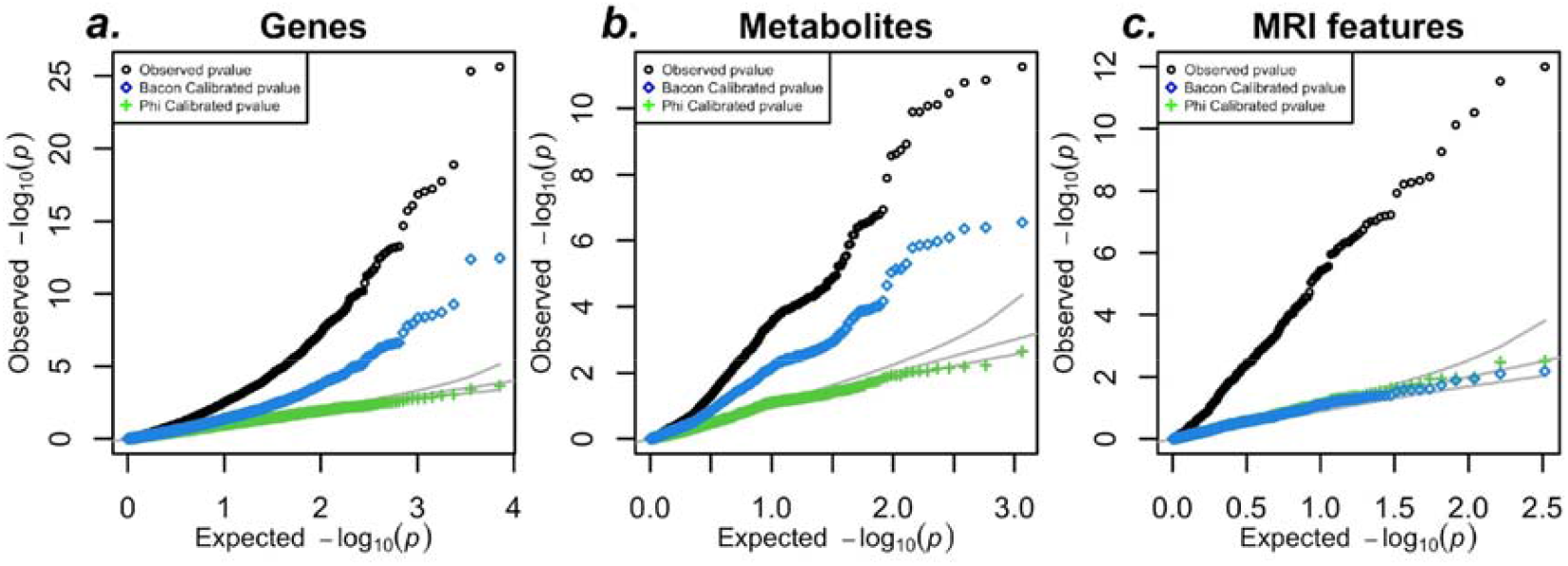
Our variance control approach corrects inflation of xWAS in UK Biobank. a) QQ-plot of the association between predicted whole blood expression and a null target trait in 100,000 UK Biobank individuals. This figure shows the results based on PredictDB models and Supplementary Figure Panel S6 shows similar inflation and correction effectiveness with Fusion models. (b) QQ-plot of the association between predicted metabolites and a null target trait in 100,000 UK Biobank individuals. (c) QQ-plot of the association between predicted MRI-derived features and a null target trait in 100,000 UK Biobank individuals. In all panels, uncorrected p-values (black) show substantial inflation. The BACON method reduces inflation but does not fully correct it, while our variance control method (green) provides well-calibrated p-values, closely aligning with the expected distribution.

### Application to actual GWAS

To facilitate the implementation of the variance control correction, we updated the S-PrediXcan software (available at https://github.com/hakyimlab/MetaXcan) to automatically apply the correction using the inflation slope parameter **Φ**. These parameters were also integrated into the database of gene expression predictors (accessible at https://predictdb.org). These enhancements simplify the correction process for end users, who only need to provide the GWAS sample size and the heritability of the target trait. GWAS sample sizes are usually available with the study and the heritability can be easily estimated from summary statistics using linkage disequilibrium score regression (LDSC) or similar methods (Bulik-Sullivan et al., 2015).

We used these updates to run a variance-controlled TWAS for three published GWAS traits: type 2 diabetes, schizophrenia, and chronic kidney disease. For illustration, we used whole blood gene expression prediction models.

As shown in Figure 6, several genes that initially appeared significant without correction—such as *PEAK1* for diabetes, *TRIM10* for schizophrenia, and *CCDC57* for chronic kidney disease—fell below the Bonferroni significance threshold after correction, underscoring the importance of this adjustment. Before correction, we identified 5, 90, and 4 Bonferroni-significant genes for diabetes, schizophrenia, and chronic kidney disease, respectively. After applying variance control to adjust for inflation, these numbers were reduced to 2 significant genes for each trait. We also performed the analysis using Fusion models and software, as shown in Supplementary Figure S7; the results were consistent with those from PredictDB, as expected.

**Figure 6:**
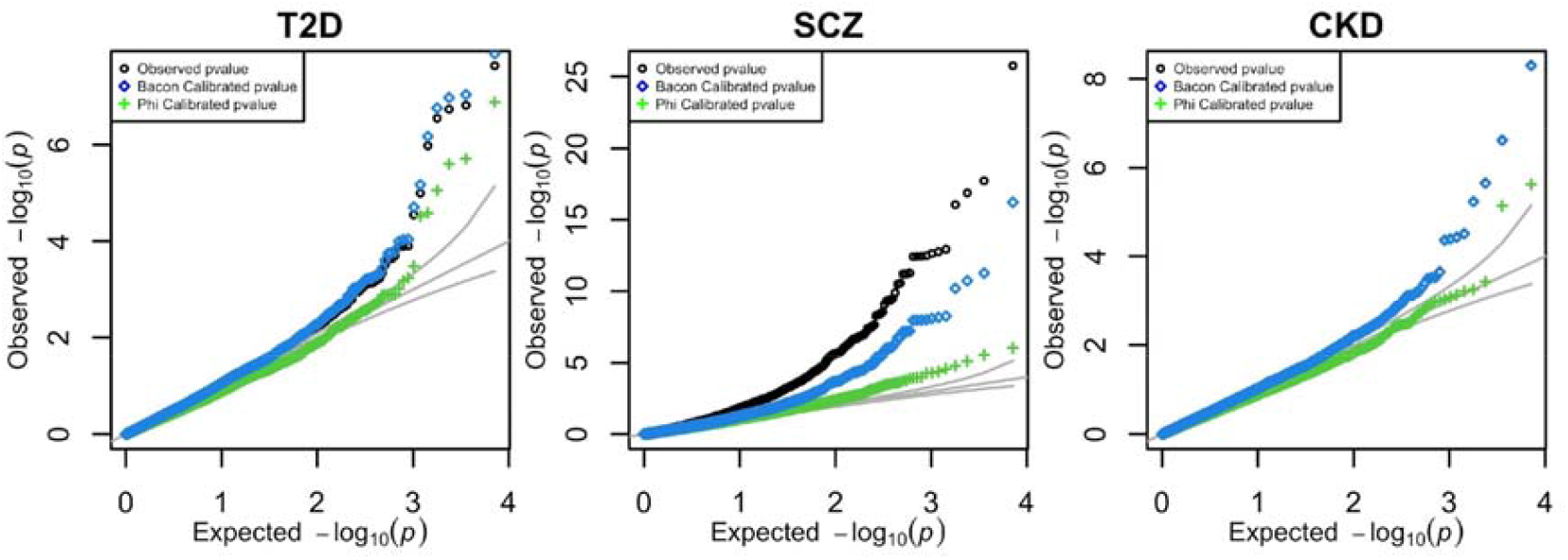
TWAS of real GWAS. Raw and corrected TWAS results for T2D, Schizophrenia, and Chronic Kidney Disease using whole blood prediction models. These results can be found in the Supplementary Table 1.

## Discussion

We reported the problem of inflation of type I error (false positive rate) in TWAS and other xWAS methods when the target traits are highly polygenic. We showed that polygenicity of the target trait causes inflated type I error both in a toy example and in a real TWAS using UK Biobank data. Given the pervasive polygenicity of most complex traits, correction of this effect is critical. We provide a user-friendly variance control approach to correct for this inflation.

The inflation is not exclusive to any one implementation of TWAS, but applies to the entire class of methods that correlate genetic predictors of gene expression—or other mediating traits—and a complex trait that is fully polygenic. Any TWAS-related method, including PrediXcan (Gamazon et al., 2015), Fusion (Gusev et al., 2016), PWAS (Brandes et al., 2020), UTMOST (Hu et al., 2019), and many others (Bhattacharya et al., 2023; Baca et al., 2022; Grishin and Gusev, 2022a), will yield an inflated false positive rate with current GWAS sample sizes if uncorrected. Analyses that correlate PRS of biomarkers or other traits with a highly polygenic target trait will suffer from the same inflation problems.

We demonstrated that inflation occurs across a variety of genetic architectures for the mediating trait. This includes gene expression, metabolite levels, and MRI-derived brain phenotypes, which range from highly sparse (i.e., gene expression) to fully polygenic (i.e., brain features). While all these traits are affected by inflation, more sparse traits have more variable inflation parameters and can reach higher values than more polygenic ones.

We proposed an effective strategy to correct for the inflation by estimating an inflation slope parameter that is specific to the mediator and valid for a large class of target traits, as long as the target trait can be well approximated by an infinitesimal model where the contribution of a single SNP is modest. Further research may improve the inflation factor calculation when the polygenic architecture of the target trait is more complex.

We corroborated error-in-variables literature by demonstrating that error-in-prediction of gene expression does not cause inflation in type I error as long as the prediction error is independent of the target trait. This assumption of independence is reasonable given the fact that prediction training is performed in studies that are independent of the GWAS studies. If the assumption does not hold, TWAS methods should be abandoned until a solution is found. We believe that most researchers would likely view this as an extreme measure and agree that, under the null, assuming independence between prediction error and the target trait is reasonable.

Some may interpret the observed inflation as resulting from horizontal pleiotropy (i.e., variants contributing to the prediction of the mediating trait that also influence the target trait through mechanisms unrelated to the mediating trait itself). However, since this horizontal pleiotropy arises from the polygenicity of the target trait, we prefer to attribute the observed inflation directly to polygenicity.

To clarify how our variance control method differs from other approaches that address horizontal pleiotropy, it’s helpful to distinguish two types of horizontal pleiotropy. In the first type, the observed association between the mediating trait and the target trait is driven by a different mediator (e.g., a different gene or protein) with a large effect size, often acting in cis. Our method does not address this type of horizontal pleiotropy, which is more effectively managed by other approaches, such as those proposed by Zhao et al. (2024); Xue et al. (2023); Mancuso et al. (2019); and Yuan et al. (2020).

Our approach instead addresses the horizontal pleiotropy that stems from the extensive polygenicity of the target trait. In a fully polygenic trait, every variant has an impact, so any variant predicting the mediator will also affect the target trait—even if the mediator itself is unrelated to the target. In this form of pleiotropy, the effects on the trait arise through a complex network of downstream processes and are generally small in magnitude, roughly proportional to 1/sqrt(M), where M is the effective number of polygenic causal variants for the target trait. Our correction strategy is necessary because methods designed for the first type of horizontal pleiotropy are computationally intensive and often require additional assumptions, which may not always hold. These methods may also be sensitive to LD mismatches between the GWAS study and the reference LD dataset, potentially reducing their robustness.

We updated the PrediXcan software and its database of gene expression prediction models to facilitate easy implementation of our correction method for our broad user base.

Our study has several limitations. First, we assumed an additive infinitesimal model for the target trait in both our simulations and theoretical derivation. However, in practice, traits may deviate from this model. While the estimation of the inflation factor, **Φ**, could be refined for different genetic architectures, we expect the infinitesimal model approximation provides a valuable first-order correction to the issue. Second, our correction does not account for horizontal pleiotropy where effects are larger than the polygenic background assumed here. Co-regulation of multiple genes by the same variants and LD contamination are also not addressed by our method. While other approaches exist to tackle these issues, they come with their own set of assumptions (Zhao et al., 2024; Mancuso et al., 2019; Yuan et al., 2020). Since each method has its own advantages and limitations, we believe our polygenicity-corrected results should be considered as part of a broader set of analyses to draw more reliable conclusions about the function of GWAS loci. Finally, our theoretical derivations were based on a linear regression framework, whereas many GWAS studies use logistic regression. Since linear regression provides a good approximation for logistic regression when the case-control ratio is balanced, we expect our results to be broadly applicable to balanced designs. However, for unbalanced designs, our method will need to be modified.

## Acknowledgement

This research has been conducted using the UK Biobank Resource under Application Number 89052.

This research used resources of the Argonne Leadership Computing Facility, which is a DOE O ce of Science User Facility supported under Contract DE-AC02-06CH11357.

This work was completed in part with resources provided by the University of Chicago’s Research Computing Center and Beagle3.

We also acknowledge resources from the Center for Research Informatics, funded by the Biological Sciences Division at the University of Chicago, with additional funding provided by the Institute for Translational Medicine, CTSA grant number 2U54TR002389-06 from the National Institutes of Health.

We thank Sarah Sumner for help editing the paper.

The following grants provided partial support to this project: R01AA029688, P30DK020595, 3R01CA242929-04S1.

## Data and code availability

Fusion prediction models can be downloaded from http://gusevlab.org/projects/fusion/. PredictDB prediction models can be downloaded from http://predictdb.org. The UK Biobank genotype data was obtained from https://www.ukbiobank.ac.uk/ under application number 89052. The code used to perform the analysis is available in GitHub: https://github.com/hakyimlab/twas-inflation. The updated MetaXcan software (v0.8.0) is available on GitHub:

https://github.com/hakyimlab/MetaXcan/releases/tag/v0.8.0 and Zenodo: https://doi.org/10.5281/zenodo.14113421.

## References

S. C. Baca, C. Singler, S. Zacharia, J.-H. Seo, T. Morova, F. Hach, Y. Ding, T. Schwarz, C.-C. F. Huang, J. Anderson, A. P. Fay, C. Kalita, S. Groha, M. M. Pomerantz, V. Wang, S. Linder, C. J. Sweeney, W. Zwart, N. A. Lack, B. Pasaniuc, D. Y. Takeda, A. Gusev, and M. L. Freedman. Genetic determinants of chromatin reveal prostate cancer risk mediated by context-dependent gene regulation. Nature Genetics, 54(9):1364–1375, Sept. 2022. ISSN 1546-1718. doi: 10.1038/s41588-022-01168-y.

A. N. Barbeira, R. Bonazzola, E. R. Gamazon, Y. Liang, Y. Park, S. Kim-Hellmuth, G. Wang, Z. Jiang, D. Zhou, F. Hormozdiari, B. Liu, A. Rao, A. R. Hamel, M. D. Pividori, F. Aguet, L. Bastarache, D. M. Jordan, M. Verbanck, R. Do, M. Stephens, K. Ardlie, M. McCarthy, S. B. Montgomery, A. V. Segrè, C. D. Brown, T. Lappalainen, X. Wen, and H. K. Im. Exploiting the GTEx resources to decipher the mechanisms at GWAS loci. Genome Biol, 22(1):49, Dec. 2021. ISSN 1474-760X. doi: 10.1186/s13059-020-02252-4. URL https://genomebiology.biomedcentral.com/articles/10.1186/s13059-020-02252-4.

A. Bhattacharya, D. D. Vo, C. Jops, M. Kim, C. Wen, J. L. Hervoso, B. Pasaniuc, and M. J. Gandal. Isoform-level transcriptome-wide association uncovers genetic risk mechanisms for neuropsychiatric disorders in the human brain. Nature Genetics, 55(12):2117–2128, Dec. 2023. ISSN 1546-1718. doi: 10.1038/s41588-023-01560-2.

E. A. Boyle, Y. I. Li, and J. K. Pritchard. An Expanded View of Complex Traits: From Polygenic to Omnigenic. Cell, 169(7):1177–1186, June 2017. Publisher: Elsevier.

N. Brandes, N. Linial, and M. Linial. Pwas: proteome-wide association study—linking genes and phenotypes by functional variation in proteins. Genome Biology, 21(1), July 2020. ISSN 1474-760X. doi: 10.1186/s13059-020-02089-x. URL http://dx.doi.org/10.1186/s13059-020-02089-x.

B. Bulik-Sullivan, H. K. Finucane, V. Anttila, A. Gusev, F. R. Day, P.-R. Loh, L. Duncan, J. R. Perry, N. Patterson, E. B. Robinson, et al. An atlas of genetic correlations across human diseases and traits. Nature genetics, 47(11):1236, 2015.

W. Fuller. Measurement Error Models. John Wiley & Sons, 1987. ISBN 978-0-471-86187-4. URL 10.1002/9780470316665.

E. R. Gamazon, H. E. Wheeler, K. P. Shah, S. V. Mozaffari, K. Aquino-Michaels, R. J. Carroll, A. E. Eyler, J. C. Denny, D. L. Nicolae, N. J. Cox, et al. A gene-based association method for mapping traits using reference transcriptome data. Nature genetics, 47(9):1091, 2015.

D. Grishin and A. Gusev. Allelic imbalance of chromatin accessibility in cancer identifies candidate causal risk variants and their mechanisms. Nature Genetics, 54(6):837–849, June 2022a. ISSN 1546-1718. doi: 10.1038/s41588-022-01075-2. URL http://dx.doi.org/10.1038/s41588-022-01075-2.

D. Grishin and A. Gusev. Allelic imbalance of chromatin accessibility in cancer identifies candidate causal risk variants and their mechanisms. Nature Genetics, 54(6):837–849, June 2022b. ISSN 1546-1718. doi: 10.1038/s41588-022-01075-2.

A. Gusev, A. Ko, H. Shi, G. Bhatia, W. Chung, B. W. Penninx, R. Jansen, E. J. De Geus, D. I. Boomsma, F. A. Wright, et al. Integrative approaches for large-scale transcriptome-wide association studies. Nature genetics, 48(3):245–252, 2016.

L. Haff. An identity for the wishart distribution with applications. Journal of Multivariate Analysis, 9(4): 531–544, 1979.

Y. Hu, M. Li, Q. Lu, H. Weng, J. Wang, S. M. Zekavat, Z. Yu, B. Li, J. Gu, S. Muchnik, Y. Shi, B. W. Kunkle, S. Mukherjee, P. Natarajan, A. Naj, A. Kuzma, Y. Zhao, P. K. Crane, H. Lu, and H. Zhao. A statistical framework for cross-tissue transcriptome-wide association analysis. Nature Genetics, 51(3): 568–576, Feb. 2019. ISSN 1546-1718. doi: 10.1038/s41588-019-0345-7. URL http://dx.doi.org/10.1038/s41588-019-0345-7.

C. d. Leeuw, J. Werme, J. E. Savage, W. J. Peyrot, and D. Posthuma. On the interpretation of transcriptome-wide association studies. PLOS Genetics, 19(9):e1010921, Sept. 2023. ISSN 1553-7404. doi: 10.1371/journal.pgen.1010921. URL https://journals.plos.org/plosgenetics/article?id=10.1371/journal.pgen.1010921.

Y. Liang, O. Melia, T. J. Caroll, T. Brettin, A. Brown, and H. K. Im. BrainXcan identifies brain features associated with behavioral and psychiatric traits using large scale genetic and imaging data. preprint, Genetic and Genomic Medicine, Feb. 2022.

N. Mancuso, M. K. Freund, R. Johnson, H. Shi, G. Kichaev, A. Gusev, and B. Pasaniuc. Probabilistic fine-mapping of transcriptome-wide association studies. Nature genetics, 51(4):675–682, 2019.

L. J. O’Connor, A. P. Schoech, F. Hormozdiari, S. Gazal, N. Patterson, and A. L. Price. Extreme polygenicityof complex traits is explained by negative selection. The American Journal of Human Genetics, 105(3): 456–476, 2019.

M. van Iterson, E. W. van Zwet, B. T. Heijmans, and the BIOS Consortium. Controlling bias and inflationin epigenome- and transcriptome-wide association studies using the empirical null distribution. Genome Biology, 18(1):19, Jan. 2017. ISSN 1474-760X. doi: 10.1186/s13059-016-1131-9.

M. Wainberg, N. Sinnott-Armstrong, N. Mancuso, A. N. Barbeira, D. A. Knowles, D. Golan, R. Ermel, A. Ruusalepp, T. Quertermous, K. Hao, J. L. M. Björkegren, H. K. Im, B. Pasaniuc, M. A. Rivas, and A. Kundaje. Opportunities and challenges for transcriptome-wide association studies. Nature Genetics, 51(4):592–599, Apr. 2019. ISSN 1546-1718. doi: 10.1038/s41588-019-0385-z.

S. Weisberg. Multiple Regression. In Applied Linear Regression, pages 47–68. John Wiley & Sons, Ltd, 2005. ISBN 978-0-471-70409-6. doi: 10.1002/0471704091.ch3. URL https://onlinelibrary.wiley.com/doi/abs/10.1002/0471704091.ch3. Section: 3 eprint: https://onlinelibrary.wiley.com/doi/pdf/10.1002/0471704091.ch3.

H. Xue, X. Shen, and W. Pan. Causal inference in transcriptome-wide association studies with invalid instruments and gwas summary data. Journal of the American Statistical Association, 118(543):1525–1537, Mar. 2023. ISSN 1537-274X. doi: 10.1080/01621459.2023.2183127. URL http://dx.doi.org/10.1080/01621459.2023.2183127.

J. Yang, B. Benyamin, B. P. McEvoy, S. Gordon, A. K. Henders, D. R. Nyholt, P. A. Madden, A. C. Heath, N. G. Martin, G. W. Montgomery, et al. Common snps explain a large proportion of the heritability for human height. Nature genetics, 42(7):565–569, 2010.

X. Yin, L. S. Chan, D. Bose, A. U. Jackson, P. VandeHaar, A. E. Locke, C. Fuchsberger, H. M. Stringham, R. Welch, K. Yu, L. Fernandes Silva, S. K. Service, D. Zhang, E. C. Hector, E. Young, L. Ganel, I. Das, H. Abel, M. R. Erdos, L. L. Bonnycastle, J. Kuusisto, N. O. Stitziel, I. M. Hall, G. R. Wagner, S. Ripatti, A. Palotie, J. Kang, J. Morrison, C. F. Burant, F. S. Collins, S. Ripatti, A. Palotie, N. B. Freimer, K. L. Mohlke, L. J. Scott, X. Wen, E. B. Fauman, M. Laakso, and M. Boehnke. Genome-wide association studies of metabolites in finnish men identify disease-relevant loci. Nature Communications, 13(1), Mar. 2022. ISSN 2041-1723. doi: 10.1038/s41467-022-29143-5. URL http://dx.doi.org/10.1038/s41467-022-29143-5.

Z. Yuan, H. Zhu, P. Zeng, S. Yang, S. Sun, C. Yang, J. Liu, and X. Zhou. Testing and controlling for horizontal pleiotropy with probabilistic Mendelian randomization in transcriptome-wide association studies. Nature Communications, 11(1):3861, July 2020. ISSN 2041-1723. doi: 10.1038/s41467-020-17668-6.

J. Zhang, D. Dutta, A. Köttgen, A. Tin, P. Schlosser, M. E. Grams, B. Harvey, CKDGen Consortium, B. Yu, E. Boerwinkle, J. Coresh, and N. Chatterjee. Plasma proteome analyses in individuals of European and African ancestry identify cis-pQTLs and models for proteome-wide association studies. Nat Genet, 54(5):593–602, May 2022. ISSN 1061-4036, 1546-1718. doi: 10.1038/s41588-022-01051-w. URL https://www.nature.com/articles/s41588-022-01051-w.

S. Zhao, W. Crouse, S. Qian, K. Luo, M. Stephens, and X. He. Adjusting for genetic confounders in transcriptome-wide association studies improves discovery of risk genes of complex traits. Nature Genetics, 56(2):336–347, Feb. 2024. ISSN 1546-1718. doi: 10.1038/s41588-023-01648-9.

Z. Zhu, F. Zhang, H. Hu, A. Bakshi, M. R. Robinson, J. E. Powell, G. W. Montgomery, M. E. Goddard, N. R. Wray, P. M. Visscher, and J. Yang. Integration of summary data from GWAS and eQTL studies predicts complex trait gene targets. Nature Genetics, 48(5):481–487, May 2016. ISSN 1546-1718. doi:10.1038/ng.3538.

