## Supplemental Figures and Notes for "Pervasive polygenicity of complex traits inflates false positive rates in transcriptome-wide association studies"

### 1 Supplementary Material

#### 1.1 Supplementary Figures

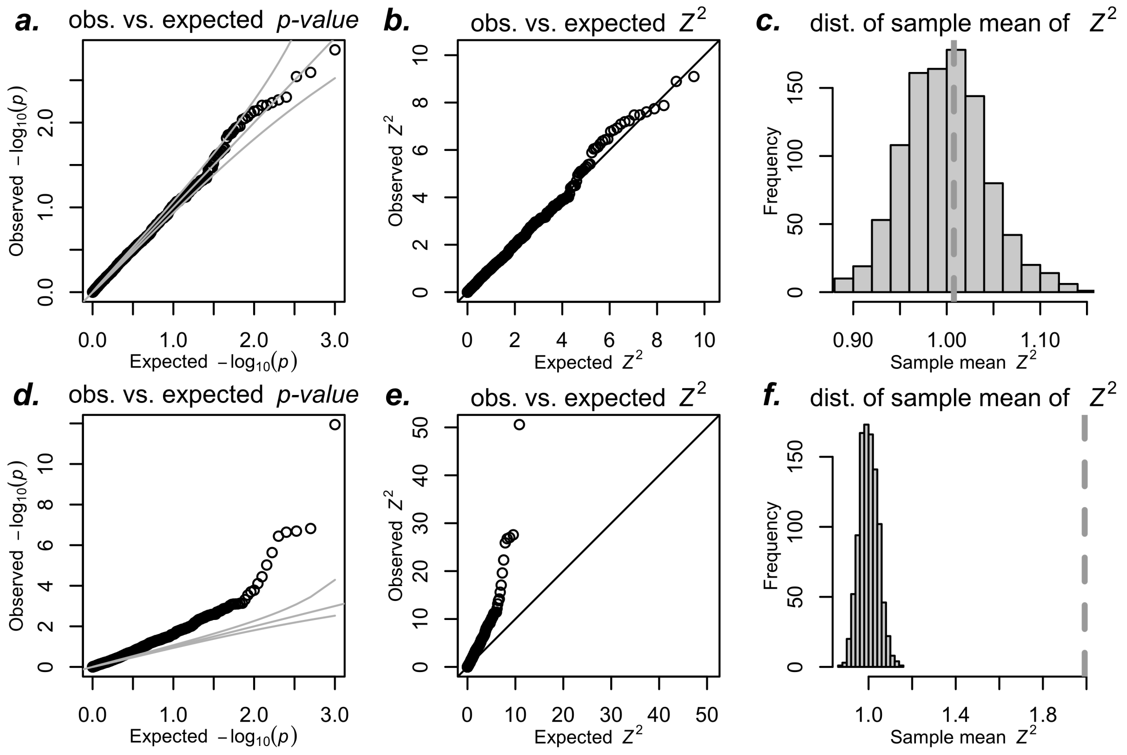


**Figure S1: Inflation in simulated TWAS - simplified setting - using student t distributed errors**. Here we simulate the $\gamma_k$, $\delta_k$, and $\epsilon_{\text{twas}}$ using the student t distribution with 2 degrees of freedom to show robustness of our results to deviations from the normal distribution in our simulations. The top row shows non-polygenic null trait simulations: (a) QQ-plot of observed p-values, (b) QQ-plot of observed $Z^2$, and (c) average $Z^2$. These follow expected distributions under the null (uniform for p-values and standard $\chi^2$ with 1 degree of freedom). The average Z 2 over 1000 simulations falls within the expected distribution of sample means of standard $\chi^2$ with 1 degree of freedom. The bottom row presents polygenic null trait simulations: (d) QQ-plot of observed p-values vs. expected, (e) QQ-plot of observed Z 2 vs expected, and (f) average Z 2 as dotted vertical line with a histogram of expected sample averages of standard normal random variables, which departs from expected distributions under the null.

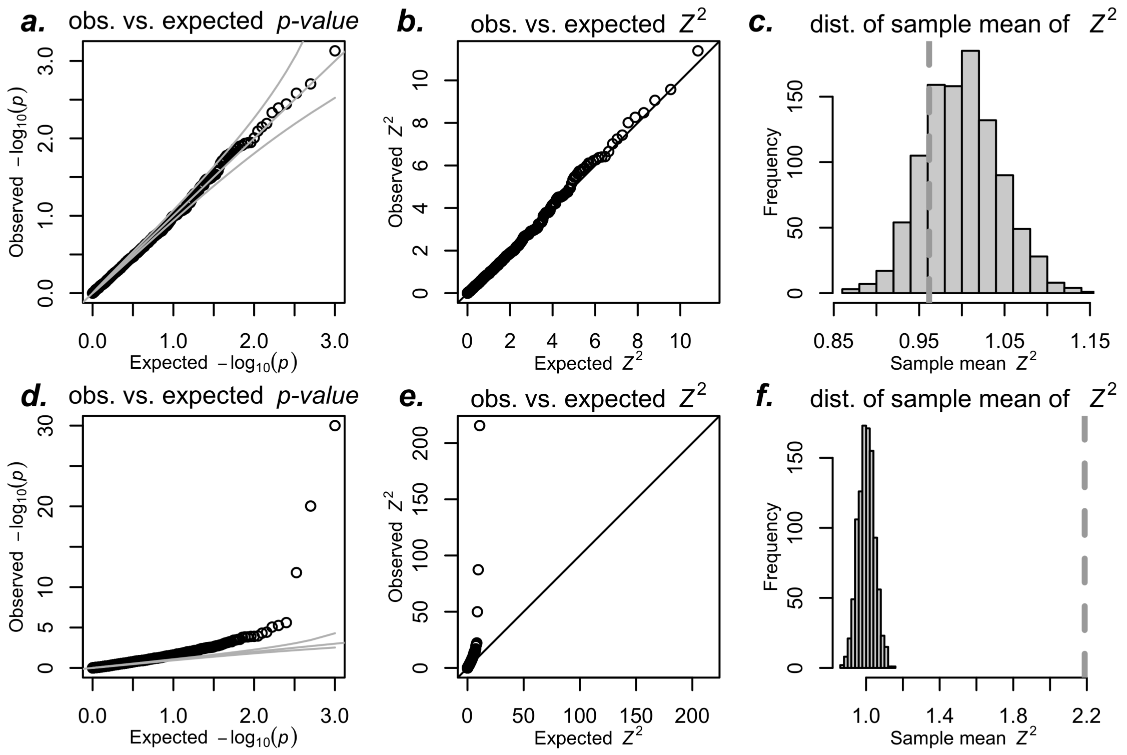


**Figure S2: Inflation in real TWAS using student t distributed errors**. We predicted the expression of the AMT gene in whole blood for 10k randomly sampled unrelated individuals from UK Biobank. For panels a-c, we simulated non-polygenic target traits from a student t distribution. For panels d-f, we simulated polygenic target traits as the sum of a polygenic component and independent student t distributed noise. We regressed the target trait on the predicted expression and calculated Z-scores, repeating this 1000 times for both trait types. Panels a and d show p-values, while b and e show squared Z-scores ($Z^2$) for non polygenic and polygenic traits, respectively. Panels c and f display the sample mean of $Z^2$ as a vertical dotted line, with histograms showing sample means of squared standard normal variables to illustrate expected means under the null.

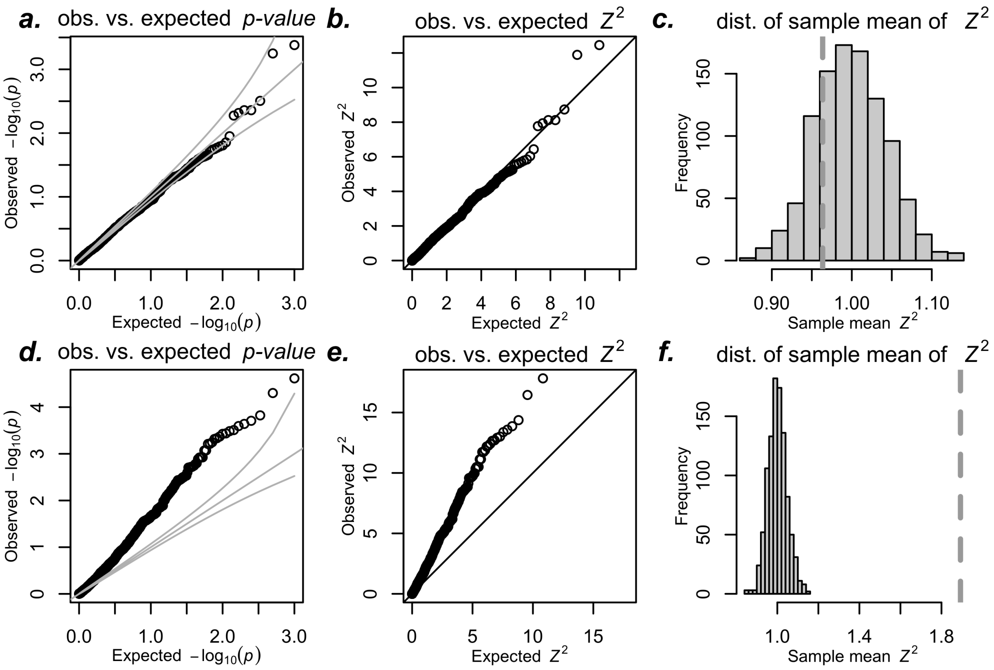


**Figure S3: Inflation in real TWAS in the UK Biobank using Fusion models**. We predicted expression of the gene AMT in whole blood for 10K randomly sampled white British unrelated individuals from the UK Biobank. For panels a-c, we simulated non-polygenic target traits from a normal distribution. For panels d-f, we simulated polygenic target traits as the sum of a polygenic component and independent normally distributed noise. We regressed the target trait on the predicted expression and calculated Z-scores, repeating this 1000 times for both trait types. Panels a and d show p-values, while b and e show squared Z-scores ($Z^2$) for non-polygenic and polygenic traits, respectively. Panels c and f display the sample mean of $Z^2$ as a vertical dotted line, with histograms showing sample means of squared standard normal variables to illustrate expected means under the null. Prediction weights for $AMT$ was downloaded from `http://gusevlab.org/projects/fusion/`.

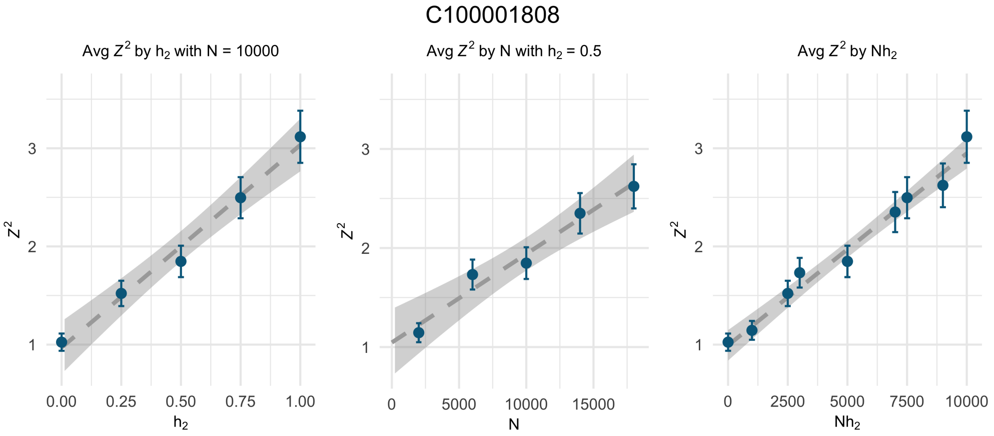


**Figure S4: Linear dependence of inflation on the GWAS sample size and heritability of the target trait**. Average $\chi^2$ (as proxy for $E\chi^2$) as functions of heritability, $h_\delta^2$, sample size, $N$, and the product of heritability and sample size, $N * h_\delta^2$. Each dot was calculated associating predicted metabolite level of *3-ethylphenylsulfate (C100001808)* with UK Biobank genotype and 1000 null polygenic traits on the same individuals at the sample sizes and heritability values indicated.

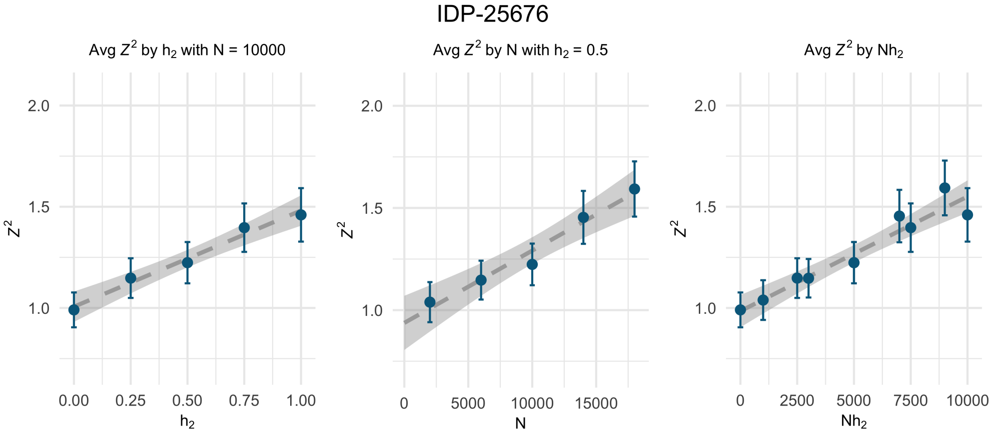


**Figure S5: Linear dependence of inflation on the GWAS sample size and heritability of the target trait**. Average $\chi^2$ (as proxy for $E\chi^2$) as functions of heritability, $h_\delta^2$, sample size, $N$, and the product of heritability and sample size, $N * h_\delta^2$. Each dot was calculated associating predicted MRI of *IDP-25676* with UK Biobank genotype and 1000 null polygenic traits on the same individuals at the sample sizes and heritability values indicated.

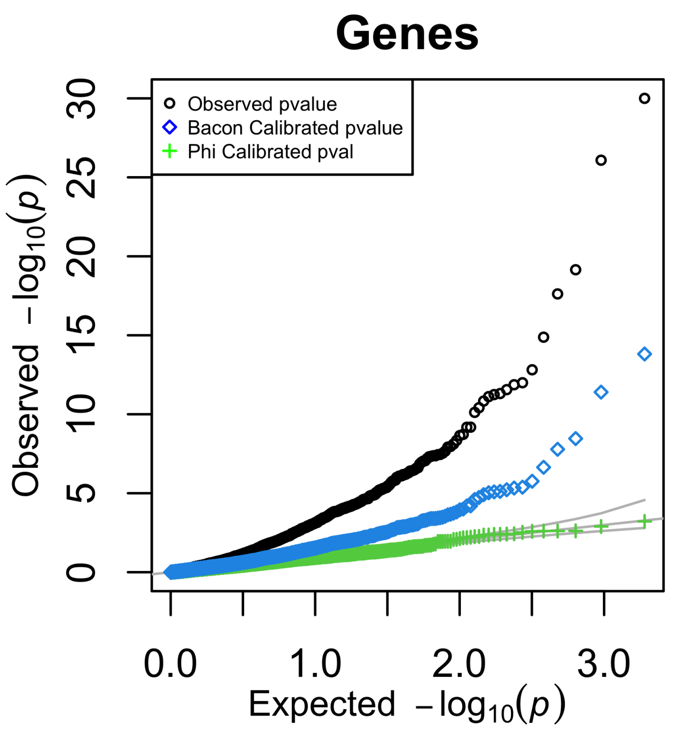


**Figure S6: Our variance control approach corrects inflation of TWAS in UK Biobank using Fusion models and software** . We simulate a null phenotype using 100k individuals from the UK Biobank and performed a GWAS to obtain the summary statistics. We use the summary statistics to run the summary based TWAS method using Fusion software to obtain association pvalues. The QQ-plot shows the pvalue distribution under the null, the uncorrected p-values are inflated (black). Bacon method reduces the inflation but fails to fully correct it. Our variance control method shown in green yields proper calibration.

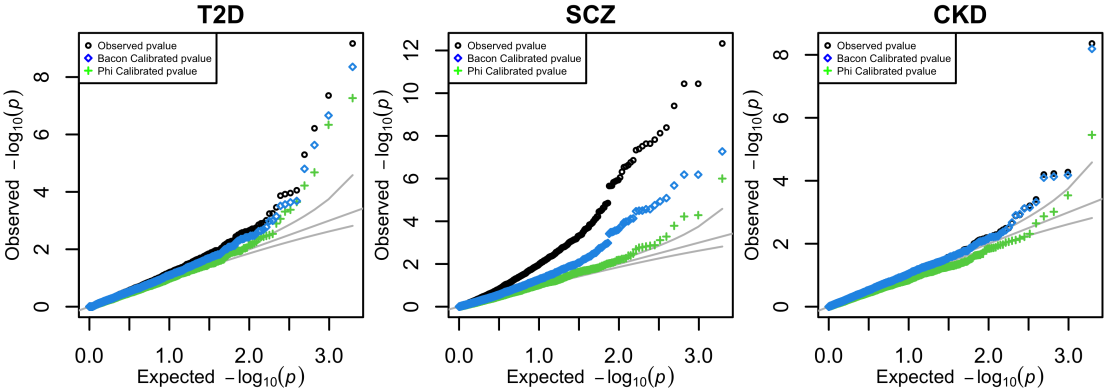


**Figure S7: Fusion TWAS of real GWAS**. Raw and corrected TWAS results for T2D, Schizophrenia, and Chronic Kidney Disease are shown.

#### 2 Supplementary Tables

| Column name | Description |
|---|---|
| gene | ensembl gene id |
| gene_name | gene symbol |
| zscore | S-PrediXcan observed gene to trait association value to show the strength and direction |
| pvalue | S-PrediXcan observed association pvalue between the gene and the trait |
| phi | individual gene phi used for calibration of TWAS results |
| corrected_pvalue | variance control method calibrated pvalue using phi, N and $h_\delta^2$ |
| bacon_corrected | bacon method calibrated pvalue |
| effect_size | S-PrediXcan's association effect size for the gene |
| pred_perf_r2 | R2 of tissue model's correlation to gene's measured transcriptome (prediction performance) |
| pred_perf_pval | pval of tissue model's correlation to gene's measured transcriptome (prediction performance) |
| pred_perf_qval | qval of tissue model's correlation to gene's measured transcriptome (prediction performance) |
| n_snps_used | number of snps from GWAS that got used in S-PrediXcan analysis |
| n_snps_in_cov | number of snps in the covariance matrix |
| n_snps_in_model | number of snps in the model |
| var_g | variance of the gene expression |
| largest_weight | weight in the prediction that has maximum magnitude |
| best_gwas_p | best snp pvalue in the gene region from the summary statistics |

**Table 1: TWAS of real GWAS traits.** We perform TWAS using the summary based method S-PrediXcan on summary statistics of 3 GWAS traits i.e Type 2 Diabetes, Schizophrenia and Chronic Kidney disease using the PredictDB whole blood models. We obtain the observed pvalue and corrected pvalues using the variance control approach and the Bacon method. The observed and corrected pvalues are show in the individual sheets for each respective traits.

### 3 Supplementary Notes on TWAS inflation

#### 3.1 TWAS model and assumptions

To examine the effect of the polygenicity of the target trait $Y$ on the association statistics, we explicitly modeled the direct genetic effects $\delta_k$ on the target trait using equation (6) below. For prediction error, we take into account that the TWAS regression is performed against a noisy version of the mediator $\tilde{T}$. Our model for the mediator, target, and noisy mediator traits are given as follows:

$$Y = \beta\, T + \sum_k X_k\ \delta_k + \epsilon, \tag{6}$$

$$T = \sum_k \gamma_k\ X_k, \text{ and} \tag{7}$$

$$\tilde{T} = \sum_k \tilde{\gamma}_k\ X_k, \tag{8}$$

where $\delta_k$, $\gamma_k$, and $\epsilon$ are continuous random variables independent of each other. $\tilde{\gamma}_k$ is assumed to be independent of $\delta_k$ and $\epsilon$. We do not assume normality for these variables but, we do assume that they have finite second moments. As in traditional TWAS, some of the $\gamma_k$'s can be 0 to include sparse architecture for the mediator. Similarly for $\tilde{\gamma}_k$'s, although we do not require that they are zero for the same set of SNPs. $k$ takes values 1 to $M$, where $M$ is the total number of causal SNPs for the trait $Y$.

To quantify the effects of target trait polygenicity and prediction error, we examined the first and second moments of the Z-score statistic of the regression of the phenotype $Y$ on the noisy predicted expression $\tilde{T}$. We call this statistic $Z_{\text{twas}}$ to emphasize the fact that this would be the results of applying standard TWAS when the polygenicity and prediction error are present but ignored.

#### 3.2 Inflation under the null

**We first show the distribution of $Z_{\text{twas}}$ under the null, i.e., $\beta = 0$.** See proof in the section 3.7. The first two moments of the Z-score are given by

$$\begin{aligned} \mathrm{E}Z_{\text{twas}} &= 0 \\ \mathrm{E}Z^2_{\text{twas}} &\approx 1 + N\ h^2_\delta\ \Phi \end{aligned} \tag{9}$$

where $N$ is the sample size of the GWAS study, and $h^2_\delta$ is the polygenic portion of $Y$, i.e., the heritability of the target trait explained by the genetic effects $\delta_k$. $\Phi$ (termed inflation slope parameter) is defined as

$$\Phi = \frac{1}{M}\ \frac{\tilde{\gamma}' \cdot \Sigma^2 \cdot \tilde{\gamma}}{\tilde{\gamma}' \cdot \Sigma \cdot \tilde{\gamma}}, \tag{10}$$

where $\Sigma$ is the limit of the LD matrix $R$ for $N \to \infty$, and $\tilde{\gamma}$ is the M-dimensional vector of estimated prediction weights in some reference training set that is typically independent of the GWAS dataset.

##### 3.2.1 Properties of the inflation slope parameter $\Phi$

The inflation in TWAS is determined by the the excess from one of the variance of the Z-score, which under the null is $N h_\delta^2 \, \Phi$. This term is positive when the target trait is polygenic, i.e., it has nonzero $h_\delta^2$. It is linear in $h_\delta^2$ and $N$. The dependence on the number and LD of the causal SNPs is encapsulated in the factor $\Phi$.

For better understanding of how $\Phi$ behaves, we investigated its properties. From the definition in equation (10), we can see that $\Phi$ is only a function of the number of causal SNPs of the target trait $M$, the prediction weights $\tilde{\gamma}$, and the LD matrix $\Sigma$, which is the large $N$ limit of $X' \cdot X/N$ and thus is no longer dependent on $N$. Therefore, $\Phi$ does not depend on the sample size, heritability, or other properties of the target trait. This indicates that $\Phi$ is a property of the mediator, which can be pre-estimated and applied to any polygenic target trait $Y$.

We also show that $\Phi$ is strictly positive, bounded below and above as follows:

$$\frac{1}{M} \leq \Phi \leq 1. \qquad (11)$$

The lower bound is attained when $\Sigma$ is the identity matrix, and the upper bound is achieved when the SNPs are perfectly correlated (See sections 3.8.3 and 3.8.4). In the oversimplified case where SNPs were independent ($\Sigma$ = identity matrix), we would have

$$\Phi = \frac{1}{M} \qquad \text{(independent SNPs)},$$

and hence $\mathrm{E}Z_{\mathrm{twas}}^2 \approx 1 + N \; (h_\delta^2/M)$ under the null.

##### 3.2.2 The factor $\Phi$ is independent of the precision of the predictor $\tau^2$

$\Phi$ is only a function of the noisy prediction weights $\tilde{\gamma}$, the LD matrix, and the total number of causal variants for the target trait (in our infinitesimal model, that is all the SNPs under consideration). Note that the definition of $\Phi$ does not depend on how different $\tilde{\gamma}$ is from the true $\gamma$, and hence precision does not play a role in the determination of $\Phi$.

#### 3.3 Inflation under the alternative

**To understand the effect of polygenicity under the alternative**, we derive a more general formula for the mean and variance of the $Z_{\mathrm{twas}}$ statistic, allowing $\beta \neq 0$.

Under the alternative, we need to specify the relationship between the true and noisy mediators. We assume that the weights $\tilde{\gamma}$ of the noisy predictor $\tilde{T}$ are given by $\tilde{\gamma} = \gamma + \epsilon_{\gamma,k}$. Hence,

$$\tilde{T} = \sum_k (\gamma_k + e_{\gamma,k})\ X_k. \tag{12}$$

where $e_{\gamma,k}$ is independent of $\gamma_k$, $\delta_k$ and $\epsilon$, which will ensure that the error in prediction is independent of the $\epsilon_{\text{twas}} = \sum_k X_k\ \delta_k + \epsilon$ and of $T$.

Under the alternative, the first two moments of the Z-score are given by

$$\begin{aligned} \mathrm{E}Z_{\text{twas}} &= \beta\tau^2 \\ \mathrm{E}Z^2_{\text{twas}} &\approx 1 + \frac{Nh^2\sigma_Y^2}{\sigma_Y^2 - \tau^2\beta^2\sigma_T^2}\ \Phi + \frac{N\ \tau^2\beta^2\sigma_T^2}{\sigma_Y^2 - \tau^2\beta^2\sigma_T^2} \end{aligned} \tag{39}$$

where $N$ is the sample size, $\beta$ is the effect of the mediator $T$ on the target trait $Y$, and $h^2_\delta$ is the polygenic portion of $Y$, i.e., the heritability of the target trait explained by the genetic effects $\delta_k$. $\Phi$ is the same as defined under the null Eq. (10). $\tau^2$ is the precision of the prediction of the mediator, i.e., the signal to noise ratio of $T$:

$$\tau^2 = \frac{\text{var}(T)}{\text{var}(\tilde{T})}. \tag{13}$$

The precision $\tau^2$ is also known as the reliability ratio in the error-in-variables literature (Fuller, 1987).

##### 3.4 Prediction error has no effect on the inflation under the null

Next, we examined the effect of the precision of the prediction of the mediator on the inflation. When the error in prediction is independent of the target trait, using $T$ or $T +$ error does not change the derivation and hence the expected $Z^2_{\text{twas}}$ should not change. Indeed, we corroborated this by verifying that, under the null, the simulated expected $Z^2$ is constant across all values of the precision as shown in Figure S8a.

Under the alternative, we find that $\mathrm{E}Z^2_{\text{twas}}$ increases monotonically with the precision of the prediction as shown in Figure S8b, indicating that prediction error reduces the power of the test as expected given the known attenuation bias effect when right hand side variables are noisy versions of the true values . Details of the simulation are described in the following section.

Consistent with established literature (Fuller, 1987), our results show that prediction error in the mediator causes a loss of power, but it does not affect the inflation under the null—contrary to the conclusion in Leeuw et al. (2023).

##### 3.5 Validation of inflation equation under the null with simulations

To assess how well this approximation works under the null, we simulated both the target trait $Y$ and the mediating trait $T$ from infinitesimal models with independent effect sizes according to the equations (6) to

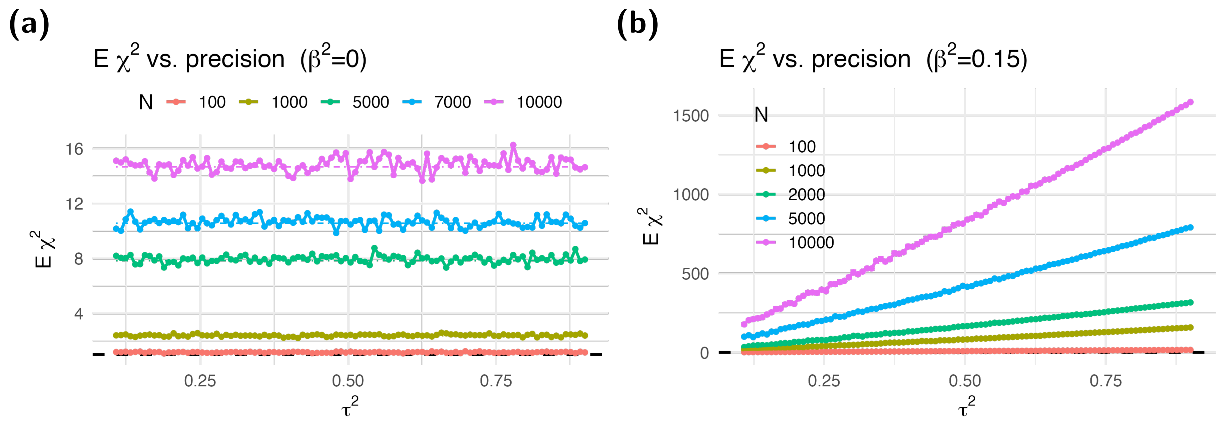


**Figure S8: Dependence of var($Z_{\text{twas}}$) on the precision $\tau^2$ of the prediction under the null and alternative.** We calculated average $Z^2$ statistics from simulations with different heritability values $h^2_\delta$ of the target trait, number of causal SNPs $M$, and sample sizes $N$. We took the average over 1,000 simulations, as well as over $h^2_\delta$ and $M$. Dash-dotted lines indicate the predicted $Z^2$ statistics from equation (9) under the null and equation (39) under the alternative.

(12). We simulated genotype data using independent binomial random variables with probability of 0.4—corresponding to the minor allele frequency of the SNPs—and therefore assumed no LD between SNPs. We used a range of values for the heritability of Y ($h^2_\delta : 0.1 - 0.9$), sample sizes ($N : 100 - 10,000$), and number of causal SNPs ($M : 99 - 6000$).

For each combination of $h^2_\delta$, $N$, and $M$, we simulated 1,000 target traits $Y$ and 99 mediating traits $T$s and $\tilde{T}$s unrelated to $Y$. Each of the 99 predicted mediating traits were simulated with different levels of precision ($\tau^2 : 0.10 - 0.9$). We then regressed each target trait on each mediating trait separately and averaged the square of the Z-scores across the 1,000 simulations, thereby obtaining an estimated $\mathrm{E}Z^2_{\text{twas}}$ for each combination of $h^2_\delta$, $N$, $M$, $T_k$, and $\tau^2$.

This estimated value was well approximated by our theoretical expression under the independent SNP assumption as shown in Figure S9a, where data points fall in the vicinity of the identity line. Panels b–d of the figure corroborate the linear relationship between the expected $Z^2_{\text{twas}}$ and $1/M$, $h^2_\delta$, and $N$, respectively, as predicted by our equation (9).

#### 3.6 Validation of inflation equation under the alternative with simulations

To examine the effect of the prediction error on the test under the alternative, we used the same simulation setup we used for the null hypothesis with $\beta^2 > 0$. Hence, we simulated the target trait $Y = \beta\, T + \sum_k \delta_k\, X_k + \epsilon$ and the mediating trait $T = \sum_k \gamma_k\, X_k$. We performed the association using the noisy version of the mediator,

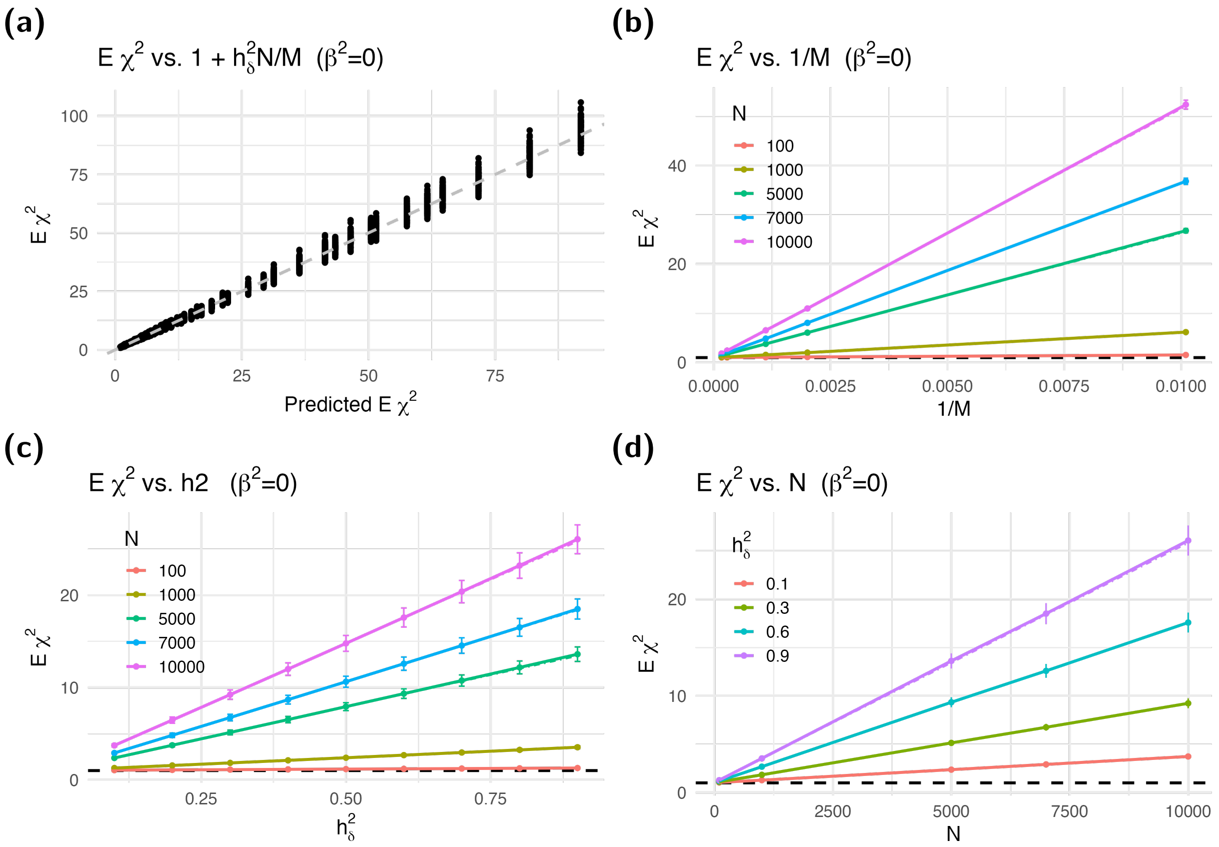


**Figure S9: Expected $Z^2$ vs. heritability, sample size, and inverse of the number of causal SNPs.** We calculated the expected association $Z^2$ statistics from simulations with different number of causal SNPs $M$, heritability values $h^2_\delta$, and sample sizes $N$. We took the average over 1,000 simulations and 99 independent mediators $T$. Panel **(a)** shows the average $Z^2$ vs $1 + h^2_\delta N/M$. . Panels **(b-d)** show the average $Z^2$ against $1/M$, $h^2_\delta$, and $N$. The horizontal dashed line at 1 indicates where calibrated $Z^2$ statistics should be. The error bars represent the 95% confidence intervals from the simulations. Dash-dotted lines in the figure show the predicted $Z^2$ statistics from equation (9) under the null—the expected $Z^2$ mostly obscures the dash-dotted line, indicating the linear relationship is consistent with the theoretical approximation. We used $M - 1$ for $M$, which further improved the match.

$\tilde{T}_k = \sum_k \tilde{\gamma}_k \ X_k$.

Similar to the null case, the estimated $\mathrm{E}Z^2_{\mathrm{twas}}$ was well approximated by our theoretical expression under the independent SNP assumption as shown in Figure S10a, where the data points fall in the vicinity of the identity line. Panels b–d of the figure corroborate the linear relationship between the expected $Z^2$ and $1/M$, $h^2_\delta$, and $N$, respectively, as predicted by our equation (29).

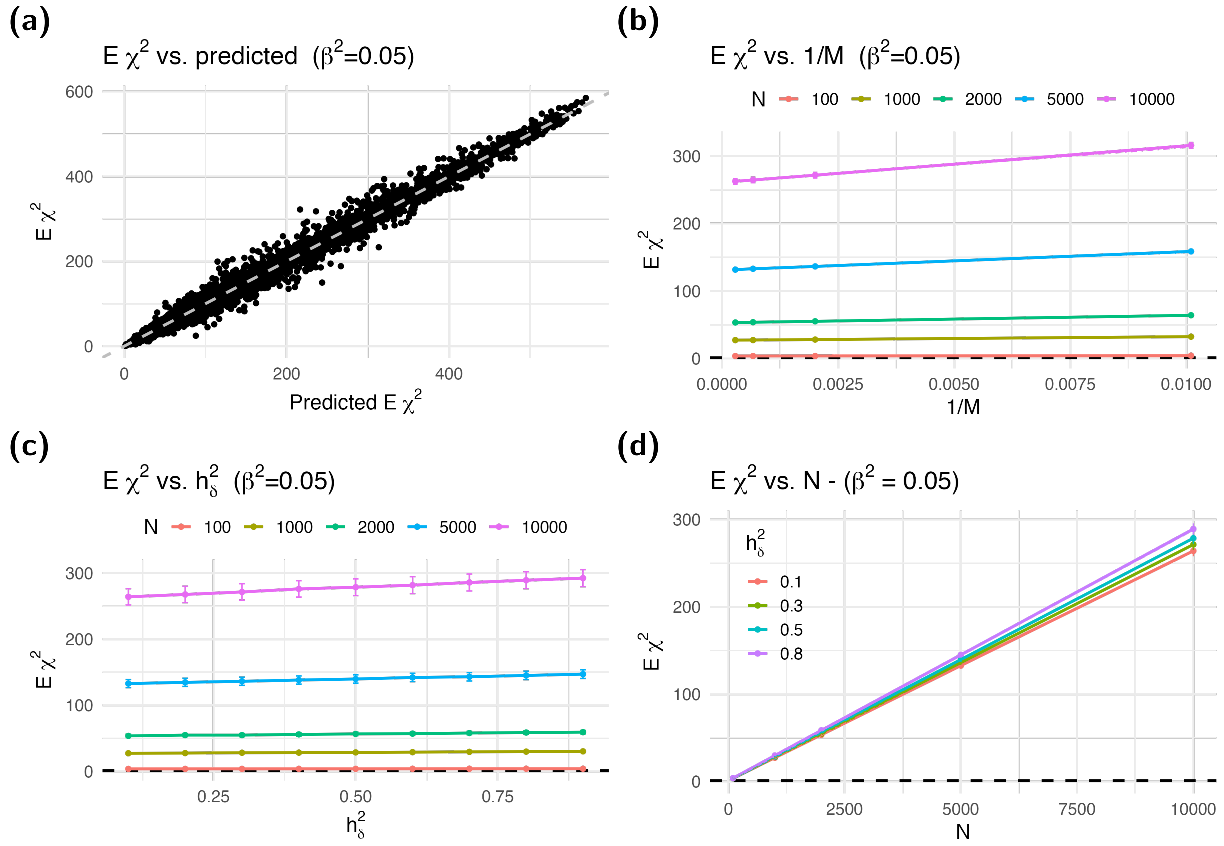


**Figure S10: Expected $Z^2$ under the alternative.** We calculated expected $Z^2$ statistics by averaging 1,000 simulations of target trait $Y$ with different combinations of number of causal SNPs $M$, polygenic portion of the heritability of $Y$ $h^2_\delta$, and sample sizes $N$. Panel **(a)** shows the expected $Z^2$ against the theoretically predicted value in equation (39) assuming independent SNPs. Dashed gray line shows the identity line. Panel **(b-d)** show the expected $Z^2$ against $1/M$, $h^2_\delta$, and $N$. Dash-dotted lines in the figure show the predicted $\mathrm{E}Z^2$ statistics from equation (39) assuming independent SNPs—for most conditions, the match is good enough to obscure the dash-dotted line in the figure. We used $M - 1$ for $M$, which further improved the match.

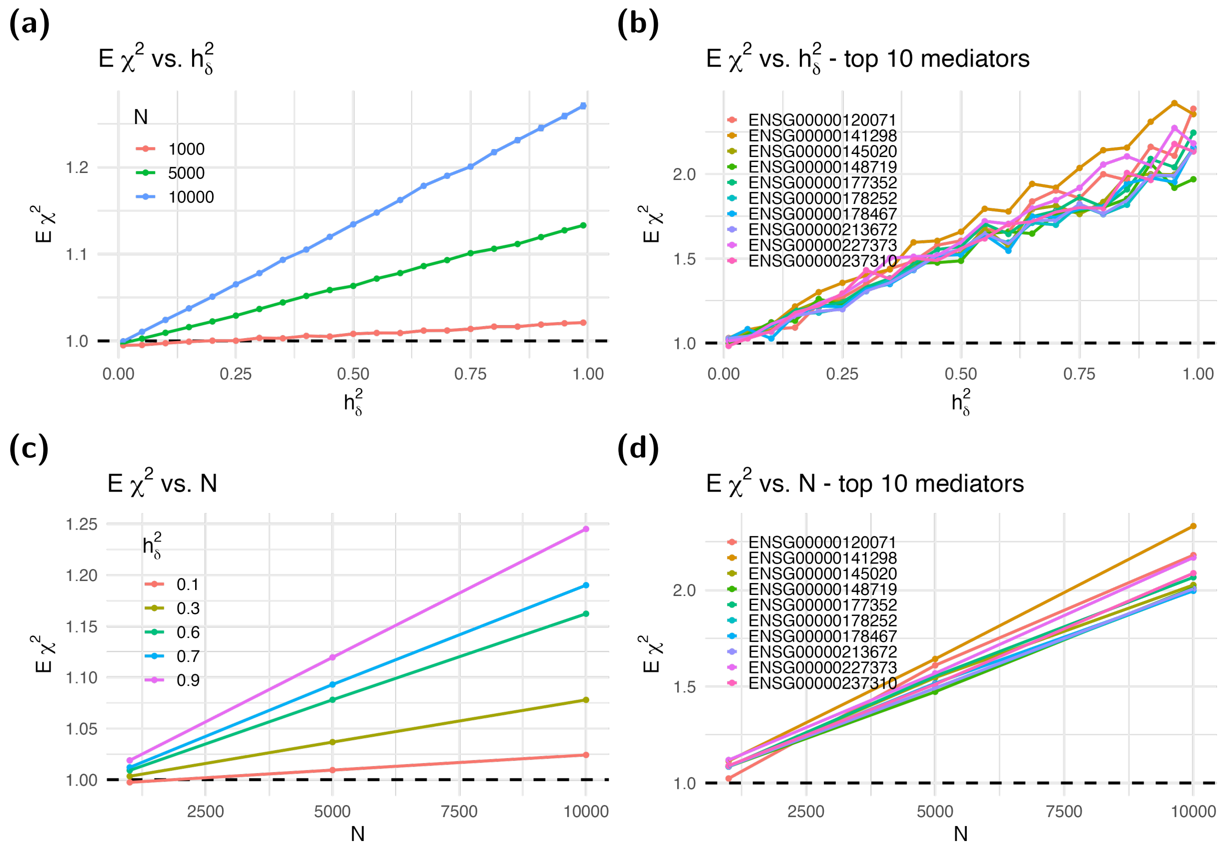


**Figure S11: Expected $Z^2$ in TWAS.** We calculated the expected $Z^2$ statistics in TWAS using simulated target traits with different heritability values $h^2_\delta$ of the target trait and sample sizes $N$. We averaged over 1,000 simulated $Y$ for a given gene, heritability, and sample size to estimate the $\mathrm{E}Z^2_{\mathrm{twas}}$ . Panel **(a)** shows the expected $Z^2$ averaged over genes as a function of the heritability $h^2_\delta$, with different lines representing different sample sizes $N$. Panel **(b)** shows the expected $Z^2$ statistics for each gene against the heritability $h^2_\delta$ of the target trait averaged over different sample sizes. Colored lines in this panel represent selected genes; we highlight the 10 genes with the highest inflation. The lines in this panel are closely linear as predicted by the formula in equation (29). Panel **(c)** shows the expected $Z^2$ against the sample size $N$, with different colors showing different heritability values $h^2_\delta$. Panel **(d)** shows the $Z^2$ statistics for each gene against the sample size $N$ averaged over different heritability values $h^2_\delta$. The lines in this panel are also closely linear, consistent with the formula in equation (29).

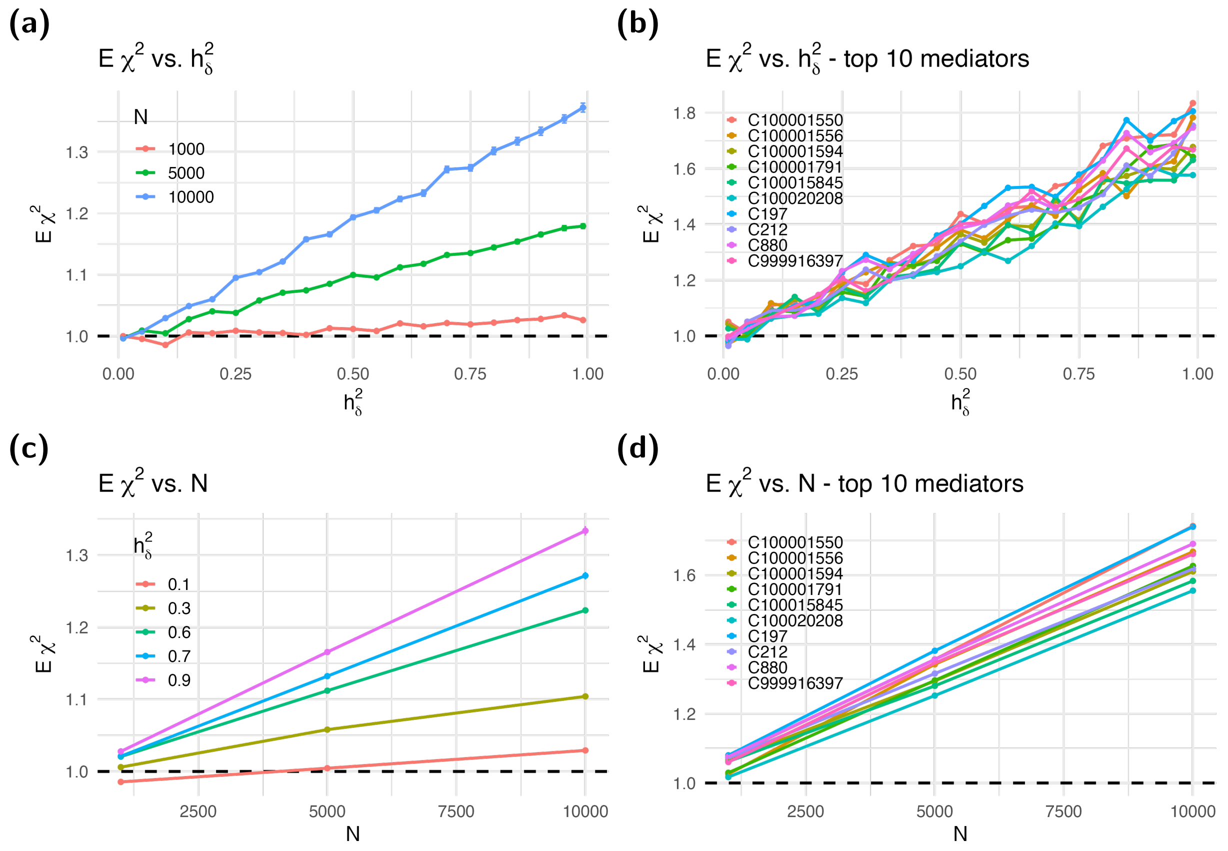


**Figure S12: Inflation in MetaboXcan.** Expected $Z^2$ statistics in TWAS calculated with simulated target traits with different heritability values of the target trait $h^2_\delta$ and sample sizes $N$. Estimated $Z^2$ is calculated averaging over 1,000 simulated $Y$ for a given metabolite, heritability, and sample size. **a** shows the average $\mathrm{E}Z^2$ over 1,156 metabolites $Z^2$ vs $h^2_\delta$. **b)** shows the $\mathrm{E}Z^2$ for each of the top 20 most inflated metabolites vs h2. **c)** shows the average $\mathrm{E}Z^2$ over 1,156 metabolites vs $N$. **d)** shows the $\mathrm{E}Z^2$ for each metabolites vs N.

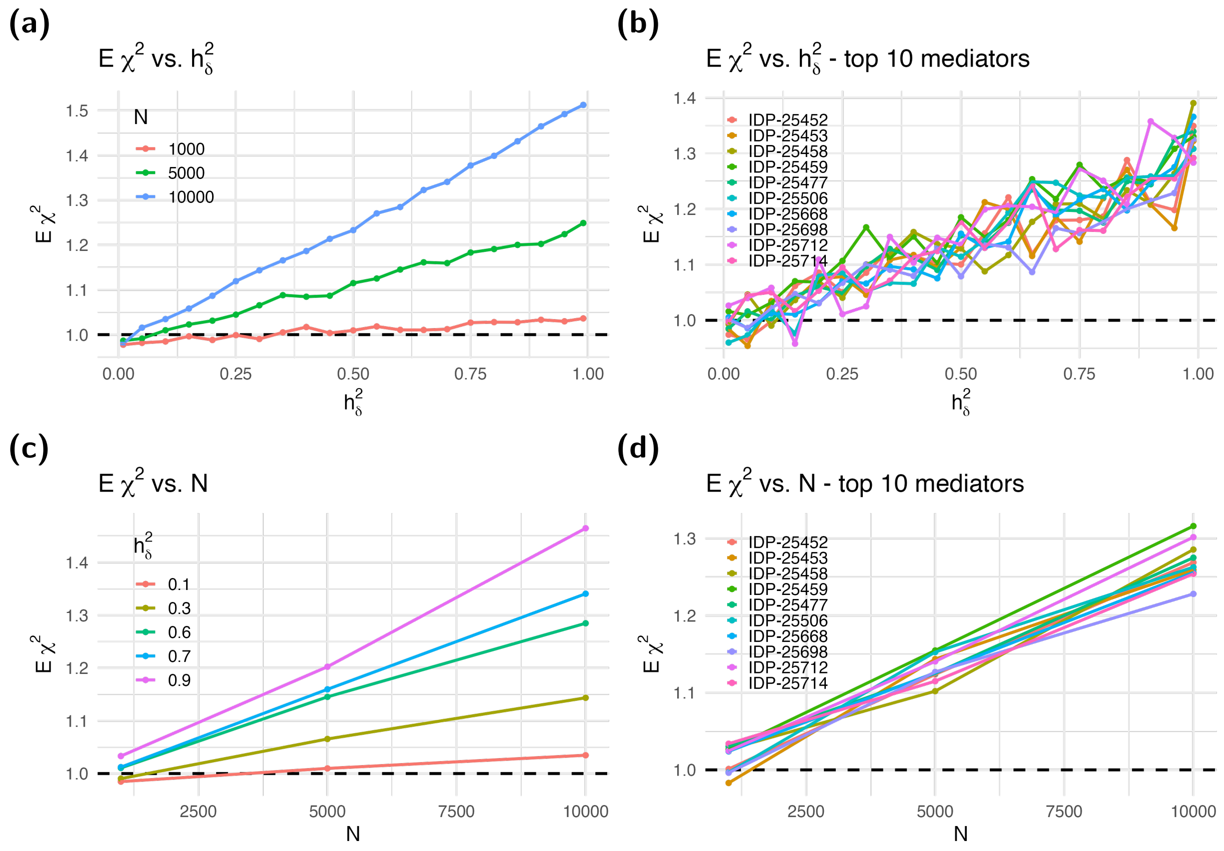


**Figure S13: Inflation in BrainXcan.** Expected $Z^2$ statistics in TWAS calculated with simulated target traits with different heritability values of the target trait $h^2_\delta$ and sample sizes $N$. Estimated $Z^2$ is calculated averaging over 1,000 simulated $Y$ for a given brain feature, heritability, and sample size. **a** shows the average $\mathrm{E}Z^2$ over 308 brain features $Z^2$ vs $h^2_\delta$. **b)** shows the $\mathrm{E}Z^2$ for each of the top 20 most inflated brain features vs h2. **c)** shows the average $\mathrm{E}Z^2$ over 308 brain features vs $N$. **d)** shows the $\mathrm{E}Z^2$ for each brain feature vs N.

##### 3.6.1 Observed inflation in TWAS with actual genotype data

To assess the practical relevance of this inflation in a traditional TWAS, we computed the association statistic between actual predicted expression levels and null polygenic target traits. We used genotype data from unrelated white British individuals in the UK Biobank with sample sizes of 1,000, 5,000 and 10,000. We predicted expression levels of 7,131 genes in whole blood using the GTEx v8 prediction models (Barbeira et al., 2021). To generate null polygenic target traits, we simulated $Y$ with heritability ranging from 0.01 to 0.99 using the same UK Biobank genotype data. We sampled the effect sizes $\delta_k$ and independent error term $\epsilon$ from independent standard normal distributions. For each combination of gene, heritability value, and sample size, we generated 1,000 independent simulated traits. We regressed out the first five genetic principal components from the simulated trait to avoid capturing associations due to population structure, which we found to be sufficient to account for population structure in our simulations. Finally, we regressed the residuals of the simulated traits against predicted expression levels and estimated the expected $Z^2_{\text{twas}}$ statistics, averaging the results across the 1,000 simulations.

Figure S11 shows the resulting average $Z^2_{\text{twas}}$ , which show a linear dependence on the heritability of the target trait and the sample size of the association consistent with equation (29). Panels (a) and (c) show the average $Z^2_{\text{twas}}$ for all genes, whereas panels (b) and (d) show the average $Z^2_{\text{twas}}$ for the top 10 genes with the highest inflation. Similar results were obtained for other mediators such as metabolites and brain features (Figure S12 and S13), showing the robustness of equation (29) to mediators with differing genetic architecture.

#### 3.7 Derivation of the distribution of the $Z_{\mathbf{twas}}$ under the null and alternative

In this section, we derive the formula for the first two moments of $Z_{\text{twas}}$

To simplify notation and derivation, we use the vector form of the model.

$$Y = \beta\, T + X \cdot \delta + \epsilon \tag{14}$$

$$T = X \cdot \gamma \tag{15}$$

$$\tilde{T} = T + X \cdot e_\gamma = T + E_T = X \cdot \tilde{\gamma} \tag{16}$$

where $\delta$, $\gamma$, $e_\gamma$, and $\epsilon$ are continuous random variables with finite second moments. $\tilde{\gamma}$ is defined as $\gamma + e_\gamma$. $\gamma$, $\delta$ are $M$-dimensional vectors with elements $\gamma_k$, $\delta_k$. $X$ is $N \times M$ genotype matrix, and $e_\gamma$ is $M$-dimensional vector. $Y$, $T$, $\tilde{T}$, $\epsilon$, and $E_T$ are $N \times 1$ vectors. Let $M$ be the number of causal SNPs for target trait $Y$, $N$ be the sample size, and $h^2_\beta$ be the heritability of the target trait explained by the mediator $T$. Clearly, from the definition above:

$$E_T = X \cdot e_\gamma \quad \text{and}$$
$$\tilde{\gamma} = \gamma + e_\gamma$$

For convenience and following standard in the field (Bulik-Sullivan et al., 2015), we use the normalized genotype matrix with mean zero and variance one.

We define the sample ($M \times M$) LD matrix as

$$R_N = R := \frac{X' \cdot X}{N}$$

and its large $N$ limit $\Sigma$ as

$$\Sigma = \lim_{N \to \infty} R \tag{17}$$

We list below the assumptions and definitions to be used for the derivation.

**TWAS assumptions and definitions**

- $\epsilon, \gamma$ continuous random variables with finite second moment
- $\epsilon, \gamma$ independent ($\perp\!\!\!\perp$), therefore $T = T = \sum_k \gamma_k X_k \perp\!\!\!\perp \epsilon$
- $\gamma$ can be sparse or polygenic

**Independent prediction error assumption**

- $e_\gamma \perp\!\!\!\perp \epsilon, \delta, \gamma$, hence $\tilde{T} \perp\!\!\!\perp X \cdot \delta$

**Polygenicity assumptions and definitions**

- $\delta_k, e_{\gamma,k}$ continuous random variables with finite second moment (no normality assumption)
- $\delta_k, \gamma_k, e_{\gamma,k}, \epsilon$ independent of each other, i.e., $T$ and $\tilde{T} \perp\!\!\!\perp X \cdot \delta$
- $M, N \gg 1$, hence $N-1 \approx N$, $M-1 \approx M$

To abbreviate notation, we use $\text{var}(Y) = \sigma_Y^2$, $\text{var}(T) = \sigma_T^2$, and $\text{var}(\tilde{T}) = \sigma_{\tilde{T}}^2$.

##### 3.7.1 Derive the estimator $\hat{\beta}$ as function of data and parameters

According to Weisberg (2005), the estimated $\hat{\beta}$ in a TWAS is

$$\begin{aligned}
\hat{\beta} &= (\tilde{T}' \cdot \tilde{T})^{-1}\ \tilde{T}' \cdot Y \\
&= (\tilde{T}' \cdot \tilde{T})^{-1} \left[\tilde{T}' \cdot (T\beta + X \cdot \delta + \epsilon)\right] \\
&= (\tilde{T}' \cdot \tilde{T})^{-1} \left[\tilde{T}' \cdot T\ \beta + \tilde{T}' \cdot X \cdot \delta + \tilde{T}' \cdot \epsilon\right] \\
&\approx \frac{1}{N\sigma_{\tilde{T}}^2} \left[\tilde{T}' \cdot T\ \beta + \tilde{T}' \cdot X \cdot \delta + \tilde{T}' \cdot \epsilon\right] \qquad \because N\ \sigma_{\tilde{T}}^2 \approx \tilde{T}' \cdot \tilde{T}
\end{aligned} \tag{18}$$

##### 3.7.2 Calculate inflation under the null

****Note that all expectations are conditional on $\tilde{T}$ and $X$ since they are known when the regression is performed. $\tilde{\gamma}$'s are also known since they are used for the prediction.****

###### 3.7.2.1 Calculate mean of $\hat{\beta}$ under the null

Since $\beta = 0$ under the null, the first term of equation (18) is 0, hence

$$\hat{\beta} = \frac{1}{\tilde{T}'\tilde{T}} \left[\tilde{T}' \cdot X \cdot \delta + \tilde{T}' \cdot \epsilon\right] \approx \frac{1}{N\text{var}(\tilde{T})} \left[\tilde{T}' \cdot X \cdot \delta + \tilde{T}' \cdot \epsilon\right]$$

The expected value of $\hat{\beta}$ is

$$\boxed{\text{E}\ \hat{\beta} = 0 \quad \text{under the null}} \tag{19}$$

since the expected value of the last two terms in (18) equal to zero, i.e., $\text{E}\left[\tilde{T}' \cdot X \cdot \delta + \tilde{T}' \cdot \epsilon \middle| \tilde{T}\right] = \tilde{T}' \cdot X \cdot \text{E}\ [\delta|\tilde{T}] + \tilde{T}' \cdot \text{E}\ [\epsilon|\tilde{T}] = \tilde{T}' \cdot X \cdot \text{E}\ [\delta] + \tilde{T}' \cdot \text{E}\ [\epsilon] = 0$ where we used that $\tilde{T} \perp\!\!\!\perp \delta$ and $\epsilon$ and $E\delta = 0$ and $E\epsilon = 0$.

###### 3.7.2.2 Calculate the limit of $\hat{\beta}$ for large N under the null

**For large** $N$

$$\boxed{\hat{\beta} = O_p(\frac{1}{\sqrt{N}}) \to 0 \quad \text{under the null}} \tag{20}$$

since 1) $\tilde{T}$ independent of $X \cdot \delta$, therefore $\sum_i^N \tilde{T}_i \; (\sum_k X_{i,k}\delta_k)/N = O_p(1/\sqrt{N})$ and 2) $\tilde{T}$ independent of $\epsilon$, therefore $\sum_i^N \tilde{T}_i \; \epsilon_i/N = O_p(1/\sqrt{N})$.

****Note that under the null, we only need to assume $\tilde{T} \perp\!\!\!\perp \epsilon_{\mathbf{twas}} = X \cdot \delta + \epsilon$, no assumptions about $T$ are needed since it's not used at all.**** Furthermore, we do not assume knowledge of the true $\gamma$'s nor of their errors, $e_\gamma$.

###### 3.7.2.3 Calculate the variance of $\hat{\beta}$ under the null

The variance of $\hat{\beta}$ estimated by TWAS is given by

$$\hat{\text{var}}(\hat{\beta}) = \frac{\hat{\text{var}}(\epsilon_{\text{twas}})}{\tilde{T}'\tilde{T}}$$

where the estimate of the variance of the error term $\hat{\text{var}}(\epsilon_{\text{twas}})$ is calculated as the residual sum of squares ($RSS$) divided by $N - 1$ (Weisberg, 2005).

$$\begin{aligned}
RSS &= (Y - \tilde{T}\;\hat{\beta})'(Y - \tilde{T}\;\hat{\beta}) \\
&= Y'Y - 2\hat{\beta}'\tilde{T}'Y + \hat{\beta}\tilde{T}'\tilde{T}\hat{\beta} \quad \because \hat{\beta}' = \hat{\beta} \quad \text{since it is a scalar} \\
&= Y'Y - \hat{\beta}^2\tilde{T}'\tilde{T} && \because (\hat{\beta}\;\tilde{T}'\tilde{T}) = \tilde{T}'Y \\
&= \sigma_Y^2 N - \hat{\beta}^2\sigma_{\tilde{T}}^2 N && \because \tilde{T}'\tilde{T} \approx N\;\sigma_{\tilde{T}}^2 \\
&\approx N\sigma_Y^2 && \because \hat{\beta}^2 = O_p(1/N) \text{ as shown in (20)}
\end{aligned}$$

$$\boxed{\hat{\text{var}}(\epsilon_{\text{twas}}) = \frac{RSS}{N-1} \approx \frac{N\sigma_Y^2}{N-1} \approx \sigma_Y^2 \quad \text{under the null}} \tag{21}$$

Therefore, the variance of $\hat{\beta}$ used in TWAS, which is unaware of the polygenic term $X \cdot \delta$, is

$$\begin{aligned}
\hat{\text{var}}(\hat{\beta}) &= \frac{\hat{\text{var}}(\epsilon_{\text{twas}})}{\tilde{T}'\tilde{T}} \\
&\approx \frac{\sigma_Y^2}{\tilde{T}'\tilde{T}} \\
\hat{\text{var}}(\hat{\beta}) &\approx \frac{\sigma_Y^2}{\sigma_{\tilde{T}}^2\;N}
\end{aligned}$$

**This variance, $\hat{\text{var}}(\hat{\beta})$, is not the actual variance of $\hat{\beta}$ when the target trait is polygenic. This discrepancy is the source of the inflation.**

###### 3.7.2.4 Calculate of E$Z_{\textbf{twas}}$ under the null

$$
\begin{aligned}
\mathrm{E}Z_{\text{twas}} &= \mathrm{E}(\frac{\hat{\beta}}{\sqrt{\mathrm{var}(\hat{\beta})}}) \\
&\approx \mathrm{E}(\hat{\beta}\ \frac{\sigma_{\tilde{T}}\sqrt{N}}{\sigma_Y}) \quad \text{for } N \gg 1 \\
&\approx \mathrm{E}\hat{\beta}\ (\frac{\sigma_{\tilde{T}}\sqrt{N}}{\sigma_Y})
\end{aligned}
$$

$$
\boxed{\mathrm{E}Z_{\text{twas}} \approx 0 \ \text{ when } N \gg 1 \quad \text{under the null}}
$$

###### 3.7.2.5 Calculate of $EZ^2_{\textbf{twas}}$ under the null

The $Z^2_{\text{twas}}$ is given by

$$
\begin{aligned}
Z^2_{\text{twas}} &\approx \frac{1}{\hat{\mathrm{var}}(\hat{\beta})} \quad \hat{\beta}^2 \\
&\approx \frac{N\ \sigma^2_{\tilde{T}}}{\sigma^2_Y} \quad \left(\frac{1}{N\sigma^2_{\tilde{T}}}\left[\tilde{T}'\cdot X\cdot\delta + \tilde{T}'\cdot\epsilon\right]\right)^2
\end{aligned}
$$

$$
\begin{aligned}
Z^2_{\text{twas}} &\approx \frac{1}{N\sigma^2_Y\sigma^2_{\tilde{T}}} \quad \left[\tilde{T}'\cdot X\cdot\delta + \tilde{T}'\cdot\epsilon\right]^2 \qquad \text{take expectation on both sides} \\
\mathrm{E}\ Z^2_{\text{twas}} &\approx \mathrm{E}\frac{1}{N\sigma^2_Y\sigma^2_{\tilde{T}}} \quad \left[\tilde{T}'\cdot X\cdot\delta + \tilde{T}'\cdot\epsilon\right]^2
\end{aligned}
$$

Rearranging terms, we get

$$
N\sigma^2_Y\sigma^2_{\tilde{T}}\ \ \mathrm{E}\ Z^2_{\text{twas}} = \mathrm{E}\left[\tilde{T}'\cdot X\cdot\delta + \tilde{T}'\cdot\epsilon\right]^2 \tag{22}
$$

When we expand the square of the terms between the brackets, cross term has expectation equal to 0 as shown below.

$$
\mathrm{E}\ [\tilde{T}'\cdot X\cdot\delta\cdot\tilde{T}'\cdot\epsilon] = \mathrm{E}\ \mathrm{E}\ [\tilde{T}'\cdot X\cdot\delta\cdot\tilde{T}'\cdot\epsilon|\delta] = \mathrm{E}\ [\tilde{T}'\cdot X\cdot\delta\cdot\tilde{T}'\cdot\mathrm{E}[\epsilon|\delta]] = 0 \tag{23}
$$

$$
\because \mathrm{E}(\epsilon|\delta) = \mathrm{E}(\epsilon) = 0 \quad \because \epsilon \perp\!\!\!\perp \delta
$$

Hence expectation on the right hand side of equation (22) is given by the expected value of the squared terms

$$
\begin{aligned}
(22) &= \text{E } (\tilde{T}' \cdot X \cdot \delta)^2 \; + \; \text{E } (\tilde{T}' \cdot \epsilon)^2 \\
&= \text{E } (\tilde{T}' \cdot X \cdot \delta \cdot \delta' \cdot X' \cdot \tilde{T}) + \text{E } (\tilde{T}' \cdot \epsilon \cdot \epsilon' \cdot \tilde{T}') && \because (\tilde{T}' \cdot X \cdot \delta) = (\tilde{T}' \cdot X \cdot \delta)' \\
& && \because (\tilde{T}' \cdot \epsilon) = (\tilde{T}' \cdot \epsilon)' \\
&= \tilde{T}' \cdot X \cdot \text{E } (\delta \cdot \delta') \cdot X' \cdot \tilde{T} + \tilde{T}' \cdot \text{E } (\epsilon \cdot \epsilon') \cdot \tilde{T}' && \text{using } E[\delta|\tilde{T}] = E[\delta] \text{ and } E[\epsilon|\tilde{T}] = E[\epsilon] \\
& && \because \tilde{T} \perp\!\!\!\perp \delta, \epsilon. \\
&= \tilde{T}' \cdot X \cdot \sigma_\delta^2 \mathbb{I}_M \cdot X' \cdot \tilde{T} + \tilde{T}' \cdot \sigma_\epsilon^2 \mathbb{I}_N \cdot \tilde{T} \\
&= \sigma_\delta^2 \; \tilde{\gamma}' \cdot X' \cdot X \cdot X' \cdot X \cdot \tilde{\gamma} + \sigma_\epsilon^2 \tilde{T}' \cdot \tilde{T} \\
&= N^2 \; \frac{h^2}{M} \; \tilde{\gamma}' \cdot R \cdot R \cdot \tilde{\gamma} + \sigma_\epsilon^2 \; \tilde{T}' \cdot \tilde{T} && \because \text{assume an infinitesimal model for } \delta, \\
& && \text{i.e., } \sigma_\delta^2 = \frac{h_\delta^2}{M}
\end{aligned} \tag{24}
$$

For clarity, here we restate both sides of the above equation (24)

$$
N\sigma_Y^2\sigma_{\tilde{T}}^2 \;\; \text{E } Z_{\text{twas}}^2 = N^2 \; \frac{h^2}{M} \; \tilde{\gamma}' \cdot R \cdot R \cdot \tilde{\gamma} + \sigma_\epsilon^2 \; \tilde{T}' \cdot \tilde{T} \tag{25}
$$

We divide both sides of the equation by $N\sigma_Y^2\sigma_{\tilde{T}}^2$

$$
\begin{aligned}
\text{E}Z_{\text{twas}}^2 &= \frac{1}{N\sigma_Y^2\sigma_{\tilde{T}}^2} \left[ N^2 \; \frac{h^2}{M} \; \tilde{\gamma}' \cdot R^2 \cdot \tilde{\gamma} + \sigma_\epsilon^2 \; \tilde{T}' \cdot \tilde{T} \right] \\
&= \frac{N}{\sigma_Y^2} \; \frac{h^2}{M} \cdot \frac{\tilde{\gamma}' \cdot R^2 \cdot \tilde{\gamma}}{\sigma_{\tilde{T}}^2} + \frac{\sigma_\epsilon^2}{\sigma_Y^2} \\
&= \frac{N}{\sigma_Y^2} \; \frac{h^2}{M} \cdot \frac{\tilde{\gamma}' \cdot R^2 \cdot \tilde{\gamma}}{\tilde{\gamma}' \cdot R \cdot \tilde{\gamma}} + \frac{\sigma_\epsilon^2}{\sigma_Y^2} \quad \because \tilde{\gamma}' \cdot R \cdot \tilde{\gamma} \approx \sigma_{\tilde{T}}^2 \\
&\approx \frac{N}{\sigma_Y^2} \; \frac{h^2}{M} \cdot \frac{\tilde{\gamma}' \cdot R^2 \cdot \tilde{\gamma}}{\tilde{\gamma}' \cdot R \cdot \tilde{\gamma}} + \frac{\sigma_Y^2(1-h_\delta^2)}{\sigma_Y^2} \quad \text{using } \sigma_Y^2(1-h_\delta^2) \\
&= \frac{Nh^2}{M} \cdot \frac{\tilde{\gamma}' \cdot R^2 \cdot \tilde{\gamma}}{\tilde{\gamma}' \cdot R \cdot \tilde{\gamma}} + 1 - h_\delta^2
\end{aligned}
$$

Therefore,

$$
\text{E } Z_{\text{twas}}^2 = 1 + Nh_\delta^2 \left( \frac{1}{M} \cdot \frac{\tilde{\gamma}' \cdot R^2 \cdot \tilde{\gamma}}{\tilde{\gamma}' \cdot R \cdot \tilde{\gamma}} - \frac{1}{N} \right) \tag{26}
$$

where $\Phi_R$ is defined as

$$
\Phi_R := \frac{1}{M} \; \frac{\tilde{\gamma}' \cdot R^2 \cdot \tilde{\gamma}}{\tilde{\gamma}' \cdot R \cdot \tilde{\gamma}} - \frac{1}{N} \tag{27}
$$

Using, $\Phi_R \approx \Phi$ (eq 45) where

$$\boxed{\Phi := \frac{1}{M}\ \frac{\tilde{\gamma}' \cdot \Sigma^2 \cdot \tilde{\gamma}}{\tilde{\gamma}' \cdot \Sigma \cdot \tilde{\gamma}}} \tag{28}$$

Finally, we have the following equation for the inflation under the null

$$\boxed{\mathrm{E} Z^2_{\text{twas}} \approx 1 + N h^2\ \Phi \quad \text{inflation under the null}} \tag{29}$$

###### 3.7.3 Calculate inflation under the alternative

$$\tilde{T}' \cdot T = (T + E_T)' \cdot T = T' \cdot T + T' \cdot E_T \approx N \mathrm{var}(T) + O_p(\sqrt{N}) \approx N \mathrm{var}(T) \quad \text{using } E_T \perp\!\!\!\perp T \tag{30}$$

###### 3.7.3.1 Calculate E $\hat{\beta}$ under the alternative

The expected value of $\hat{\beta}$ is (restating equation (18))

$$\begin{aligned}
\mathrm{E}\hat{\beta} &\approx \mathrm{E}\left(\frac{1}{N\mathrm{var}(\tilde{T})}\left[\tilde{T}' \cdot T\ \beta + \tilde{T}' \cdot X \cdot \delta + \tilde{T}' \cdot \epsilon\right]\right) \\
&\approx \frac{1}{N\mathrm{var}(\tilde{T})}\ \ \mathrm{E}\ \ \tilde{T}' \cdot T\ \beta + \frac{1}{N\mathrm{var}(\tilde{T})}\ \ \mathrm{E}\left[\tilde{T}' \cdot X \cdot \delta + \tilde{T}' \cdot \epsilon\right] \\
&\approx \frac{1}{N\mathrm{var}(\tilde{T})}\ \ \mathrm{E}\ \ N\mathrm{var}(T)\ \beta \quad \because (30) \text{ and } (23) \\
&\approx \frac{\mathrm{var}(T)}{\mathrm{var}(\tilde{T})}\ \beta
\end{aligned}$$

therefore with $\tau^2 := \frac{\mathrm{var}(T)}{\mathrm{var}(\tilde{T})}$, we get

$$\boxed{\mathrm{E}\ \hat{\beta} = \tau^2\ \beta} \tag{31}$$

###### 3.7.3.2 Calculate the limit of $\hat{\beta}$ for large N under the alternative

since in equation (18) $X \cdot \delta / N = O_p(1/\sqrt{N})$ and $\tilde{T}' \cdot \epsilon / N = O_p(1/\sqrt{N})$ for large $N$ using $X \perp\!\!\!\perp \delta\ \tilde{T} \perp\!\!\!\perp \epsilon$

$$\boxed{\hat{\beta} \longrightarrow \tau^2\ \beta \qquad \text{for } N \gg 1} \tag{32}$$

###### 3.7.3.3 Calculate var $\hat{\beta}$ under the alternative

From standard regression results (Weisberg, 2005), the variance of $\hat{\beta}$ is estimated as

$$\hat{\mathrm{var}}(\hat{\beta}) = \frac{\hat{\mathrm{var}}(\epsilon_{\text{twas}})}{\tilde{T}'\tilde{T}}$$

where the estimate of the variance of the error term $\hat{\text{var}}(\epsilon_{\text{twas}})$ is calculated as the residual sum of squares ($RSS$) divided by $N-1$ (Weisberg, 2005).

$$\begin{aligned}
RSS &= (Y - \tilde{T}\ \hat{\beta})'(Y - \tilde{T}\ \hat{\beta}) \\
&= Y'Y - 2\hat{\beta}'\tilde{T}'Y + \hat{\beta}'\tilde{T}'\tilde{T}\hat{\beta} \\
&= Y'Y - \hat{\beta}^2\tilde{T}'\tilde{T} \qquad\qquad \because (\hat{\beta}\ \tilde{T}'\tilde{T}) = \tilde{T}'Y \\
&\approx Y' \cdot Y - \beta^2\tau^2\tau^2\tilde{T}'\tilde{T} \\
&\approx N\sigma_Y^2 - \beta^2\tau^2\frac{T'T}{\tilde{T}'\tilde{T}}\tilde{T}'\tilde{T} \\
&= N\sigma_Y^2 - \tau^2\beta^2 T'T \\
&= N(\sigma_Y^2 - \tau^2\beta^2\sigma_T^2)
\end{aligned}$$

$$\boxed{\hat{\text{var}}(\epsilon_{\text{twas}}) = \frac{RSS}{N-1} \approx \frac{N(\sigma_Y^2 - \tau^2\beta^2\sigma_T^2)}{N-1} \approx \sigma_Y^2 - \tau^2\beta^2\sigma_T^2} \tag{33}$$

The variance of $\hat{\beta}$ used in TWAS, which is unaware of the polygenic term $X \cdot \delta$, is

$$\hat{\text{var}}_{\text{twas}}(\hat{\beta}) = \frac{\hat{\text{var}}(\epsilon_{\text{twas}})}{\tilde{T}'\tilde{T}} = \frac{\sigma_Y^2 - \tau^2\beta^2\sigma_T^2}{N\sigma_{\tilde{T}}^2}$$

###### 3.7.3.4 Calculate mean of $Z_{\text{twas}}$ under the alternative

$$\begin{aligned}
\text{E}Z_{\text{twas}} &= \text{E}(\frac{\hat{\beta}}{\sqrt{\text{var}(\hat{\beta})}}) \\
&\approx \text{E}(\hat{\beta}\ \frac{\sigma_{\tilde{T}}\sqrt{N}}{\sqrt{\sigma_Y^2 - \tau^2\beta^2\sigma_T^2}}) \quad \text{for } N \gg 1 \\
&\approx \text{E}\hat{\beta}\ (\frac{\sigma_{\tilde{T}}\sqrt{N}}{\sqrt{\sigma_Y^2 - \tau^2\beta^2\sigma_T^2}})
\end{aligned}$$

$$\boxed{\text{E}Z_{\text{twas}} \approx \tau^2\beta\frac{\sigma_{\tilde{T}}\sqrt{N}}{\sqrt{\sigma_Y^2 - \tau^2\beta^2\sigma_T^2}} \quad \text{when } N \gg 1 \quad \text{under the alternative}}$$

###### 3.7.3.5 Calculate $\text{E}Z^2_{\text{twas}}$ under the alternative

$$\begin{aligned}
Z^2_{\text{twas}} &= \frac{\hat{\beta}^2}{\hat{\text{var}}_{\text{twas}}(\hat{\beta})} \\
&\approx \hat{\beta}^2\ \frac{N\sigma_{\tilde{T}}^2}{\sigma_Y^2 - \tau^2\beta^2\sigma_T^2}
\end{aligned}$$

The $Z^2$ statistics in TWAS is given by

$$\begin{aligned}
Z^2_{\text{twas}} &= \hat{\beta}^2 \; \frac{N\sigma^2_{\tilde{T}}}{\sigma^2_Y - \tau^2\beta^2\sigma^2_T} \\
&= (\frac{1}{N\sigma^2_{\tilde{T}}})^2 \left[\tilde{T}' \cdot T \; \beta + \tilde{T}' \cdot X \cdot \delta + \tilde{T}' \cdot \epsilon\right]^2 \; \frac{N\sigma^2_{\tilde{T}}}{\sigma^2_Y - \tau^2\beta^2\sigma^2_T} \\
&= (\frac{1}{N\sigma^2_{\tilde{T}}}) \left[\tilde{T}' \cdot T \; \beta + \tilde{T}' \cdot X \cdot \delta + \tilde{T}' \cdot \epsilon\right]^2 \; \frac{1}{\sigma^2_Y - \tau^2\beta^2\sigma^2_T}
\end{aligned}$$

Take expectation on both sides

$$\text{E} \; Z^2_{\text{twas}} = \frac{1}{N\sigma^2_{\tilde{T}}} \text{E} \left[N \; \sigma^2_T \beta + \tilde{T}' \cdot X \cdot \delta + \tilde{T}' \cdot \epsilon\right]^2 \; \frac{1}{\sigma^2_Y - \tau^2\beta^2\sigma^2_T}$$

Rearranging terms, we get

$$(\sigma^2_Y - \tau^2\beta^2\sigma^2_T) \; (N\sigma^2_{\tilde{T}}) \text{E} \; Z^2_{\text{twas}} = \text{E} \left[N \; \sigma^2_T \; \beta + \tilde{T}' \cdot X \cdot \delta + \tilde{T}' \cdot \epsilon\right]^2 \tag{34}$$

When we expand the square of the terms between the brackets, all the cross terms have expectation equal to 0 as shown below.

$$\begin{aligned}
&\text{E} \; [N \; \beta \; \tilde{T}' \cdot X \cdot \delta] = N \; \beta \; \tilde{T}' \cdot X \cdot \text{E} \; [\delta] = 0 && \because \text{E} \; \delta = 0 \\
&\text{E} \; [N \; \beta \; \tilde{T}' \cdot X \cdot \epsilon] = N \; \beta \; \tilde{T}' \cdot X \cdot \text{E} \; [\epsilon] = 0 && \because \text{E} \; \epsilon = 0 \\
&\text{E} \; [\tilde{T}' \cdot X \cdot \delta \cdot \tilde{T}' \cdot \epsilon] = \text{E} \; \text{E} \; [\tilde{T}' \cdot X \cdot \delta \cdot \tilde{T}' \cdot \epsilon|\delta] = \text{E} \; [\tilde{T}' \cdot X \cdot \delta \cdot \tilde{T}' \cdot \text{E}[\epsilon|\delta]] = 0 \\
&\because \text{E}(\epsilon|\delta) = \text{E}(\epsilon) = 0 \quad \because \epsilon \perp\!\!\!\perp \delta
\end{aligned}$$

Therefore, the expectation is given by the expected value of the squared terms

$$\begin{aligned}
(34) &= \text{E} \; (N \; \sigma^2_T \; \beta)^2 \; + \; \text{E} \; (\tilde{T}' \cdot X \cdot \delta)^2 \; + \; \text{E} \; (\tilde{T}' \cdot \epsilon)^2 \\
&= N^2 \; \sigma^4_T \; \beta^2 + \text{E} \; (\tilde{T}' \cdot X \cdot \delta \cdot \delta' \cdot X' \cdot \tilde{T}) + \text{E} \; (\tilde{T}' \cdot \epsilon \cdot \epsilon' \cdot \tilde{T}') && \because (\tilde{T}' \cdot X \cdot \delta) = (\tilde{T}' \cdot X \cdot \delta)' \\
& && \because (\tilde{T}' \cdot \epsilon) = (\tilde{T}' \cdot \epsilon)' \\
&= N^2 \; \sigma^4_T \; \beta^2 + \tilde{T}' \cdot X \cdot \text{E} \; (\delta \cdot \delta') \cdot X' \cdot \tilde{T}) + \tilde{T}' \cdot \text{E} \; (\epsilon \cdot \epsilon') \cdot \tilde{T}' \\
&= N^2 \; \sigma^4_T \; \beta^2 + \tilde{T}' \cdot X \cdot \sigma^2_\delta \mathbb{I}_M \cdot X' \cdot \tilde{T} + \tilde{T}' \cdot \sigma^2_\epsilon \mathbb{I}_N \cdot \tilde{T} \\
&= N^2 \; \sigma^4_T \; \beta^2 + \sigma^2_\delta \; \tilde{\gamma}' \cdot X' \cdot X \cdot X' \cdot X \cdot \tilde{\gamma} + \sigma^2_\epsilon \tilde{T}' \cdot \tilde{T} \\
&= N^2 \; \sigma^4_T \; \beta^2 + N^2 \; \frac{\sigma^2_Y \; h^2}{M} \; \tilde{\gamma}' \cdot R \cdot R \cdot \tilde{\gamma} + \sigma^2_\epsilon \; \tilde{T}' \cdot \tilde{T} && \because \text{assume an infinitesimal model for } \delta, \\
& && \text{i.e., } \sigma^2_\delta = \frac{\sigma^2_Y \; h^2_\delta}{M} \\
&= N^2 \; \sigma^4_T \; \beta^2 + N^2 \; \frac{\sigma^2_Y \; h^2}{M} \; \tilde{\gamma}' \cdot R^2 \cdot \tilde{\gamma} + \sigma^2_\epsilon \; N \; \sigma^2_{\tilde{T}}
\end{aligned} \tag{35}$$

For clarity, here we restate both sides of the above equation (35)

$$(\sigma^2_Y - \tau^2\beta^2\sigma^2_T) \; (N\sigma^2_{\tilde{T}}) \; \text{E} \; Z^2_{\text{twas}} = N^2 \; \sigma^4_T \; \beta^2 + N^2 \; h^2 \; \frac{\tilde{\gamma}' \cdot R^2 \cdot \tilde{\gamma}}{M} + \sigma^2_\epsilon \; N \; \sigma^2_{\tilde{T}} \tag{36}$$

We divide both sides of the equation by $N\ \sigma_{\tilde{T}}^2$

$$
\begin{aligned}
(\sigma_Y^2 - \tau^2\beta^2\sigma_T^2)\ \mathrm{E}Z_{\text{twas}}^2 &= \frac{1}{N\ \sigma_{\tilde{T}}^2}\left[N^2\ \sigma_T^4\ \beta^2 + N^2\ \frac{\sigma_Y^2\ h^2}{M}\ \tilde{\gamma}' \cdot R^2 \cdot \tilde{\gamma} + \sigma_\epsilon^2\ N\sigma_{\tilde{T}}^2\right] \\
&= N\beta^2\sigma_T^2\ \tau^2 + N\frac{\sigma_Y^2\ h^2}{M\ \sigma_{\tilde{T}}^2} \cdot \tilde{\gamma}' \cdot R^2 \cdot \tilde{\gamma} + \sigma_\epsilon^2 \\
&\approx N\beta^2\sigma_T^2\ \tau^2 + N\frac{\sigma_Y^2\ h^2}{M} \cdot \frac{\tilde{\gamma}' \cdot R^2 \cdot \tilde{\gamma}}{\tilde{\gamma}' \cdot R \cdot \tilde{\gamma}} + \sigma_\epsilon^2 \quad \text{using } \tilde{\gamma}' \cdot R \cdot \tilde{\gamma} \approx \sigma_{\tilde{T}}^2 \\
&= N\beta^2\sigma_T^2\ \tau^2 + N\frac{\sigma_Y^2\ h^2}{M} \cdot \frac{\tilde{\gamma}' \cdot R^2 \cdot \tilde{\gamma}}{\tilde{\gamma}' \cdot R \cdot \tilde{\gamma}} + \sigma_Y^2 - \beta^2\sigma_T^2 - h_\delta^2\sigma_Y^2 \\
&= \sigma_Y^2 - \tau^2\beta^2\sigma_T^2 + \tau^2\beta^2\sigma_T^2 + N\beta^2\sigma_T^2\ \tau^2 + N\frac{\sigma_Y^2\ h^2}{M} \cdot \frac{\tilde{\gamma}' \cdot R^2 \cdot \tilde{\gamma}}{\tilde{\gamma}' \cdot R \cdot \tilde{\gamma}} - \beta^2\sigma_T^2 - h_\delta^2\sigma_Y^2 \\
&= (\sigma_Y^2 - \tau^2\beta^2\sigma_T^2) + \left(\tau^2\beta^2\sigma_T^2 + N\ \tau^2\beta^2\sigma_T^2 - \beta^2\sigma_T^2\right) + \\
&\quad N\sigma_Y^2\ h^2\left(\frac{1}{M}\ \frac{\tilde{\gamma}' \cdot R^2 \cdot \tilde{\gamma}}{\tilde{\gamma}' \cdot R \cdot \tilde{\gamma}} - 1/N\right) \\
&= \left(\sigma_Y^2 - \tau^2\beta^2\sigma_T^2\right) + N\ \tau^2\beta^2\sigma_T^2\left(\frac{1}{N} + 1 - \frac{1}{N\tau^2}\right) + N\sigma_Y^2\ h^2\left(\frac{1}{M}\ \frac{\tilde{\gamma}' \cdot R^2 \cdot \tilde{\gamma}}{\tilde{\gamma}' \cdot R \cdot \tilde{\gamma}} - 1/N\right) \\
&\approx \left(\sigma_Y^2 - \tau^2\beta^2\sigma_T^2\right) + N\ \tau^2\beta^2\sigma_T^2 + N\sigma_Y^2\ h^2\left(\frac{1}{M}\ \frac{\tilde{\gamma}' \cdot R^2 \cdot \tilde{\gamma}}{\tilde{\gamma}' \cdot R \cdot \tilde{\gamma}} - 1/N\right)
\end{aligned}
$$

The last approximation uses, $1 + \frac{1}{N} - \frac{1}{N\tau^2} \approx 1$.

Dividing both sides by $\left(\sigma_Y^2 - \tau^2\beta^2\sigma_T^2\right)$, we get

$$\mathrm{E}Z_{\text{twas}}^2 \approx 1 + \frac{Nh^2\sigma_Y^2}{\sigma_Y^2 - \tau^2\beta^2\sigma_T^2}\ \Phi_R + \frac{N\ \tau^2\beta^2\sigma_T^2}{\sigma_Y^2 - \tau^2\beta^2\sigma_T^2}$$

where $\Phi_R$ is defined as

$$\Phi_R := \frac{1}{M}\ \frac{\tilde{\gamma}' \cdot R^2 \cdot \tilde{\gamma}}{\tilde{\gamma}' \cdot R \cdot \tilde{\gamma}} - \frac{1}{N} \tag{37}$$

Using, $\Phi_R \approx \Phi$ (eq 45) where

$$\boxed{\Phi := \frac{1}{M}\ \frac{\tilde{\gamma}' \cdot \Sigma^2 \cdot \tilde{\gamma}}{\tilde{\gamma}' \cdot \Sigma \cdot \tilde{\gamma}}} \tag{38}$$

So, for general prediction weights $\tilde{\gamma}$,

$$\boxed{\mathrm{E}Z_{\text{twas}}^2 \approx 1 + \frac{Nh^2\sigma_Y^2}{\sigma_Y^2 - \tau^2\beta^2\sigma_T^2}\ \Phi + \frac{N\ \tau^2\beta^2\sigma_T^2}{\sigma_Y^2 - \tau^2\beta^2\sigma_T^2} \qquad \text{general inflation formula}} \tag{39}$$

#### 3.8 Additional proofs and derivation of intermediate results

##### 3.8.1 Proof that $\mathbf{E}_R(R^2) = \Sigma^2 + \frac{\Sigma^2 + \mathbf{tr}(\Sigma)\Sigma}{N}$

(Note that we are using $\mathrm{E}_R$ to indicate that $R$, a sample covariance matrix, is being integrated out, not conditioned on as in the rest of the derivation. Here we are not taking expectation with respect to $R$ but using the fact that $R$ and its quadratic forms are converging to their expected values.)

From Theorem 3.1 in (Haff, 1979), for a random variable $S \sim W_M(\Sigma, N)$,

$$\mathrm{cov}(S_{ij}, S_{kl}) = N \cdot (\Sigma_{ik}\Sigma_{jl} + \Sigma_{il}\Sigma_{jk})$$

Recall that $N \cdot R \sim W_M(\Sigma, N)$ so $R = \frac{S}{N}$.

$$\begin{aligned}
[\ \mathrm{E}_R(R^2)\ ]_{ij} &= \mathrm{E}_R(R_i' R_j) \\
&= \mathrm{E}_R(\sum_k R_{ki} R_{kj}) \\
&= \sum_k \mathrm{E}_R(R_{ki} R_{kj}) \qquad (40) \\
\mathrm{E}_R(R_{ki} R_{kj}) &= \mathrm{E}_R(R_{ki})\mathrm{E}_R(R_{kj}) + \mathrm{cov}(R_{ki}, R_{kj}) \\
&= \Sigma_{ki}\Sigma_{kj} + \frac{1}{N^2} \cdot N \cdot (\Sigma_{kk}\Sigma_{ij} + \Sigma_{ki}\Sigma_{kj}) \\
&= \Sigma_{ki}\Sigma_{kj} + \frac{\Sigma_{kk}\Sigma_{ij} + \Sigma_{ki}\Sigma_{kj}}{N} \qquad (41) \\
\therefore\ \mathrm{E}_R(R^2) &= \Sigma^2 + \frac{\Sigma^2 + \mathrm{tr}(\Sigma)\Sigma}{N}, \quad \text{from (40) and (41)} \qquad (42)
\end{aligned}$$

##### 3.8.2 Proof that $\Phi_R \approx \frac{1}{M}\ \frac{\tilde{\gamma}' \cdot \Sigma^2 \cdot \tilde{\gamma}}{\tilde{\gamma}' \cdot \Sigma \cdot \tilde{\gamma}} =: \Phi$ and using this for the final version of $\Phi$

We can approximate the following expression using the asymptotic approximations of $R^2$ and $R$ since we are interested in large $N$ settings

$$\frac{\tilde{\gamma}' \cdot R^2 \cdot \tilde{\gamma}}{\tilde{\gamma}' \cdot R \cdot \tilde{\gamma}}$$

As stated in Equation (17), $R \approx \Sigma$ for large $N$. With the genotype of $i$th sample being an M-dimensional column vector $X_{i\cdot}$,

$$\begin{aligned}
R^2 &= \frac{\sum_i X_{i\cdot} X_{i\cdot}'}{N} \cdot \frac{\sum_j X_{j\cdot} X_{j\cdot}'}{N} \\
&= \frac{1}{N^2} \sum_{i,j} X_{i\cdot} X_{i\cdot}' X_{j\cdot} X_{j\cdot}' \\
&\approx \mathrm{E}_R[\ R^2\ ] \quad \because \text{ law of large numbers and } \mathrm{E} X_{i\cdot} X_{i\cdot}' X_{j\cdot} X_{j\cdot}' = \mathrm{E}_R[\ R^2\ ] \qquad (43)
\end{aligned}$$

Here we use $\mathrm{E}_R$ to indicate that we are integrating over genotype $X$'s. With Equations (17) and (43), we have the following approximation:

$$\begin{aligned}
\frac{\tilde{\gamma}' \cdot R^2 \cdot \tilde{\gamma}}{\tilde{\gamma}' \cdot R \cdot \tilde{\gamma}} &\approx \frac{\tilde{\gamma}' \cdot \mathrm{E}_R[\ R^2\ ] \cdot \tilde{\gamma}}{\tilde{\gamma}' \cdot \Sigma \cdot \tilde{\gamma}} \\
&= \frac{\tilde{\gamma}' \cdot (\Sigma^2 + \frac{\Sigma^2 + \mathrm{tr}(\Sigma)\Sigma}{N}) \cdot \tilde{\gamma}}{\tilde{\gamma}' \cdot \Sigma \cdot \tilde{\gamma}} \quad \text{, see (42)} \\
&\approx \frac{\tilde{\gamma}' \cdot (\Sigma^2 + \frac{M}{N}\Sigma) \cdot \tilde{\gamma}}{\tilde{\gamma}' \cdot \Sigma \cdot \tilde{\gamma}} \\
&= \frac{\tilde{\gamma}' \cdot \Sigma^2 \cdot \tilde{\gamma}}{\tilde{\gamma}' \cdot \Sigma \cdot \tilde{\gamma}} + \frac{M}{N} \qquad (44)
\end{aligned}$$

From equations (37) and (44), we have

$$\begin{aligned}
\Phi_R &\approx \frac{1}{M} \cdot \left( \frac{\tilde{\gamma}' \cdot \Sigma^2 \cdot \tilde{\gamma}}{\tilde{\gamma}' \cdot \Sigma \cdot \tilde{\gamma}} + \frac{M}{N} \right) - \frac{1}{N} \\
&= \frac{1}{M} \cdot \left( \frac{\tilde{\gamma}' \cdot \Sigma^2 \cdot \tilde{\gamma}}{\tilde{\gamma}' \cdot \Sigma \cdot \tilde{\gamma}} \right) \qquad (45)
\end{aligned}$$

###### 3.8.3 Proof that $1/M \leq \Phi$

Recall

$$\Phi = \frac{1}{M} \frac{\tilde{\gamma}' \cdot \Sigma^2 \cdot \tilde{\gamma}}{\tilde{\gamma}' \cdot \Sigma \cdot \tilde{\gamma}}$$

$$\sigma^2_{\tilde{T}} = \tilde{\gamma}' \cdot R \cdot \tilde{\gamma} \approx \tilde{\gamma}' \cdot \Sigma \cdot \tilde{\gamma},$$

To find an extreme of $\Phi$ , we will find an extreme of

$$\tilde{\gamma}' \cdot \Sigma^2 \cdot \tilde{\gamma}$$

with the condition that

$$\sigma^2_{\tilde{T}} = \tilde{\gamma}' \cdot \Sigma \cdot \tilde{\gamma},$$

using the Lagrange multiplier approach, i.e., we will find the extreme of the function

$$f(\lambda, \mathcal{L}) = \tilde{\gamma}' \cdot \Sigma^2 \cdot \tilde{\gamma} - \mathcal{L}\ (\tilde{\gamma}' \cdot \Sigma \cdot \tilde{\gamma} - \sigma^2_{\tilde{T}}),$$

where $\mathcal{L}$ is the Lagrange multiplier. We will also write the numerator in a more convenient form by using the eigenvalue decomposition of $\Sigma$. Since $\Sigma$ is symmetric, it can be eigenvalue-decomposed

$$\Sigma = C \cdot \Lambda \cdot C'$$

where $C$ is the matrix of eigenvectors of $\Sigma$ as columns and is orthonormal $C'C = \mathbb{I}$. $\Lambda$ is a diagonal matrix with the eigenvalues of $\Sigma$, $\lambda_k$'s.

$$\begin{aligned}\tilde{\gamma}' \cdot \Sigma \cdot \tilde{\gamma} &= \tilde{\gamma}' \cdot C \cdot \Lambda \cdot C' \cdot \tilde{\gamma} \\ &= \gamma_C' \cdot \Lambda \cdot \gamma_C \quad \text{with } \gamma_C := C' \cdot \tilde{\gamma} \\ &= \sum_k \gamma_{C,k}^2 \ \lambda_k\end{aligned}$$

Since the eigenvalues of $\Sigma^2$ are the squares of the eigenvalues of $\Sigma$, we have

$$\begin{aligned}\tilde{\gamma}' \cdot \Sigma^2 \cdot \tilde{\gamma} &= \tilde{\gamma}' \cdot C \cdot \Lambda^2 \cdot C' \cdot \tilde{\gamma} \\ &= \gamma_C' \cdot \Lambda^2 \cdot \gamma_C \\ &= \sum_k \gamma_{C,k}^2 \lambda_k^2\end{aligned}$$

Now, we rewrite the function $f$ in terms of the eigenvalues of $\Sigma$

$$f(\lambda, \mathcal{L}) = \sum_k \gamma_{C,k}^2 \ \lambda_k^2 - \mathcal{L} \left( \sum_k \gamma_{C,k}^2 \ \lambda_k - \sigma_{\tilde{T}}^2 \right)$$

where $\mathcal{L}$ is the Lagrange multiplier. The solution is obtained by setting the derivatives with respect to $\lambda_k$ and $\mathcal{L}$ to 0.

$$\begin{gathered}2 \ \gamma_{C,k}^2 \ \lambda_k - \mathcal{L} \ \gamma_{C,k}^2 = 0 \text{ and} \\ \sum_k \gamma_{C,k}^2 \ \lambda_k - \sigma_{\tilde{T}}^2 = 0\end{gathered}$$

Therefore,

$$\lambda_k = \frac{\mathcal{L}}{2} \quad \text{and} \quad \sum_k \gamma_{C,k}^2 \ \frac{\mathcal{L}}{2} = \sigma_{\tilde{T}}^2$$

$$\mathcal{L} = \frac{2 \ \sigma_{\tilde{T}}^2}{\sum_l \gamma_{C,l}^2} \implies \lambda_k = \frac{\sigma_{\tilde{T}}^2}{\sum_l \gamma_{C,l}^2} \ \forall \ k$$

We can show that this is indeed a minimum of the function $f$ by calculating the Hessian of the function and verifying that all the eigenvalues are positive. $f'' = 2\gamma_{C,k}^2 > 0$ for all $k$ and cross derivatives are 0, so that the Hessian is diagonal so that its eigenvalues are $2\gamma_{C,k}^2 > 0$. Therefore, this solution corresponds to a minimum.

The minimum of the function corresponds to the case where all the eigenvalues are equal; therefore, $\Sigma$ is the identity matrix.

If we plug in the identity matrix in the equation for $\Phi$, we get

$$\boxed{\frac{1}{M} \leq \Phi}$$

since $\frac{\tilde{\gamma}' \cdot \Sigma^2 \cdot \tilde{\gamma}}{\tilde{\gamma}' \cdot \Sigma \cdot \tilde{\gamma}} = \frac{\tilde{\gamma}' \tilde{\gamma}}{\tilde{\gamma}' \tilde{\gamma}} = 1$

##### 3.8.4 Proof that $\Phi \leq 1$

Recall

$$\Phi = \frac{1}{M} \frac{\tilde{\gamma}' \cdot \Sigma^2 \cdot \tilde{\gamma}}{\tilde{\gamma}' \cdot \Sigma \cdot \tilde{\gamma}}$$

Using the eigenvalue decomposition of $\Phi$, we can re-express as

$$\Phi = \frac{1}{M} \frac{\sum_k \gamma_{C,k}^2 \lambda_k^2}{\sum_k \gamma_{C,k}^2 \lambda_k}$$

Since the trace of $\Sigma$ is $M$, $\sum_k \lambda_k = M$ and $\lambda \geq 0$, $\lambda_k \leq M$

Hence using $\lambda_k^2 \leq \lambda_k \ M$

$$\Phi \leq \frac{1}{M} \frac{\sum_k \gamma_{C,k}^2 \lambda_k \ M}{\sum_k \gamma_{C,k}^2 \lambda_k} = 1$$

$$\boxed{\Phi \leq 1} \tag{46}$$

This upper bound is attained when the first eigenvalue equals M and all others are 0, i.e., all SNPs are perfectly correlated as seen below

$$\Phi = \frac{1}{M} \frac{\gamma_{C,1}^2 M^2}{\gamma_{C,1}^2 M} = 1.$$

Since variance of $\tilde{T}$ has to be non zero, $\gamma_{C,1}^2$ is $> 0$.

##### 3.8.5 Decomposition of $\sigma_\epsilon^2$

$$\sigma_\epsilon^2 = \sigma_Y^2 - \sigma_T^2 \beta^2 - h_\delta^2 \sigma_Y^2$$

since $\sigma_Y^2 = \text{var}(Y) = \text{var}(\beta T) + \text{var}(X \cdot \delta) + \text{var}(\epsilon)$, given the independence of $T$, $\epsilon$, and $\delta$

##### 3.8.6 Show that $\gamma' \cdot R \cdot \gamma \approx \sigma_T^2$

$$\begin{aligned} \sigma_T^2 &\approx \hat{\text{var}}(T) \\ &= \hat{\text{var}}(X \cdot \gamma) \\ &\approx \gamma' \cdot X' \cdot X \cdot \gamma / N \\ &= \gamma' \cdot R \cdot \gamma \end{aligned}$$

so we have

$$\boxed{\gamma' \cdot R \cdot \gamma \approx \sigma_T^2} \tag{47}$$

##### 3.8.7 Show that $\mathbf{tr}(R) = M$

$$\mathrm{tr}(R) = \sum_k R_{kk} = \sum_k \sum_i X'_{ki} X_{ik}/N = \sum_k \sum_i X_{ik}^2/N = \sum_k \hat{\mathrm{var}}(X_k) = \sum_k 1 = M \tag{48}$$

Analogous derivation for $\tilde{\gamma}$ yields

$$\tilde{\gamma}' \cdot R^2 \cdot \tilde{\gamma} = \mathrm{var}(\tilde{\gamma})\mathrm{tr}(R^2) \tag{49}$$

$$\tilde{\gamma}' \cdot R \cdot \tilde{\gamma} = \mathrm{var}(\tilde{\gamma})\mathrm{tr}(R) \tag{50}$$
